# Target-Stabilized Base Editors Enable Robust High-Fidelity RNA Editing

**DOI:** 10.1101/2025.07.31.667791

**Authors:** Taian Liu, Yunping Lin, Qiwei Liu, Wenhui Liao, Yujing Zhang, Wenhua Cao, Yunyi Lin, Lixin Yang, Zexuan Hong, Zhonghua Lu

**Affiliations:** Research Center for Primate Neuromodulation and Neuroimaging, Institute of Biomedical and Health Engineering, Shenzhen Institutes of Advanced Technology, Chinese Academy of Sciences, Shenzhen 518055, China; Shenzhen Key Laboratory for Molecular Biology of Neural Development, Shenzhen Technological Research Center for Primate Translational Medicine, Shenzhen-Hong Kong Institute of Brain Science, Shenzhen Institutes of Advanced Technology, Chinese Academy of Sciences, Shenzhen 518055, China; University of Chinese Academy of Sciences, Beijing 100049, China; Shenzhen Maternity and Child Healthcare Hospital, Southern Medical University, Shenzhen 518027, China; State Key Laboratory of Biomedical Imaging Science and System, Chinese Academy of Sciences, Shenzhen 518055, China

## Abstract

RNA base editing using engineered deaminases expands target compatibility, offering a powerful tool to correct mutations at the RNA level. However, widespread off-target effects, primarily arising from dissociated free deaminases, remain a significant challenge. Here, we devised the RECODE (<u>R</u>NA <u>e</u>diting with <u>co</u>n<u>d</u>itionally stable and <u>e</u>nhanced ADAR1 deaminase variants) system, which employs designer degron-tagged ADAR1 deaminase (ADAR1d) with guide RNA (gRNA)-regulated stability. By promoting degradation of gRNA-unbound ADAR1d, RECODE markedly reduced transcriptome-wide edits while maintaining high on-target efficacy. Engineering gRNA for target RNA-induced conformational switching restricts ADAR1d stabilization, further enhancing editing precision. With structure-guided rational engineering of ADAR1d, RECODE achieved editing efficiencies of 90.3% and 58.8% for an Amyotrophic Lateral Sclerosis-relevant *FUS* mutation *in vitro* and *in vivo*, respectively. These findings establish RECODE as a highly stringent and efficient RNA editing technology and underscore a general principle for enhancing the specificity of RNA-guided protein effectors.

## Introduction

Biological organisms rely not only on the faithful transmission of genetic information from DNA to RNA and subsequently to proteins, as described by the central dogma, but also engage in a variety of post-transcriptional processes that modify RNA molecules, including RNA sequences^1–3^. These include alternative RNA splicing^4^, 3′-end processing^5^, and RNA editing^6, 7^, which collectively enhance molecular diversity and adaptation to external stimuli. Among metazoans, adenosine-to-inosine (A-to-I) conversion—catalyzed by double-stranded RNA-specific adenosine deaminases (ADARs)—represents the most prevalent form of RNA editing^7^. This modification is crucial for functions such as immune regulation^8–10^, cancer progression^11^, and temperature adaptation^12, 13^. Since inosine is read as guanosine by the cellular machinery^14^, and G-to-A mutations are the second most common type of pathogenic single-nucleotide variants (SNVs) ^15^, ADAR-mediated RNA editing holds considerable promise for correcting disease-causing mutations.

Compared to CRISPR-based DNA editing techniques, RNA editing offers distinct advantages in clinical applications, such as reversibility and tunability, derived from the transient nature of RNA. These characteristics make RNA correction potentially safer and particularly beneficial in contexts requiring temporal pharmacodynamic control^16^. However, in contrast to the rapid development of DNA editing tools, considerably less effort has been devoted to creating programmable RNA base editing platforms.

In recent years, several strategies have emerged to direct ADAR deaminases to specific RNA sequences for site-directed RNA editing (SIDE). There are two main categories that these approaches fall into based on their reliance upon either endogenous ADARs or exogenous deaminases. In the first category, native ADAR1 and/or ADAR2 are recruited to target sites by a long double-stranded RNA (dsRNA) segment formed by target-guide hybrids^17–19^ or by structured ADAR-recruitment motifs^20–23^. While effective in several *in vivo* studies using animal models^18, 23–25^, these strategies are constrained by the poorly understood principles of ADAR substrate selectivity^26, 27^, in addition to the variable expression levels of ADARs across different tissues and cellular states^28, 29^. The use of exogenous deaminases offers a solution to these constraints, with additional benefits such as an expandable base and context compatibility^30, 31^. However, the widespread off-target effects from overexpressed deaminases^19, 23, 32, 33^ and delivery challenges, particularly for CRISPR/Cas-based effectors^34–38^, limit their potential clinical applicability. Moreover, deaminase mutagenesis for enhanced precision often results in a trade-off with overall activity^34,38^.

Here, we present a strategy termed RECODE (RNA editing with conditionally stable and enhanced ADAR1 deaminase variants), which enables selective elimination of guide RNA (gRNA)-unbound ADAR1 deaminases (ADAR1d) that cause promiscuous edits, whilst maintaining robust on-target activity. We further enhanced editing precision by engineering gRNA for target RNA-induced conformational switching. In addition, we report that RECODE demonstrated significant therapeutic potential by effectively editing an Amyotrophic Lateral Sclerosis (ALS)-relevant mutation in the *FUS* gene and mitigating FUS mislocalization to neuronal axons. Finally, we have expanded the utility of RECODE by installing protective mutations in *Angptl3* and targeting start codons for translational interference.

## Results

### A novel engineered degradation tag, UDeg3a, has an increased dynamic range of responsiveness to Pepper

To eliminate dissociated free deaminase responsible for global off-target edits, we aimed to generate a conditional deaminase that becomes unstable when it escapes from the editing complex. We employed the recently developed Pepper-tDeg system, an RNA aptamer-regulated protein degradation tag^39^. In this system, tDeg serves as a C-terminal degron that renders the protein unstable unless bound to the RNA aptamer Pepper. We envisioned that fusion with tDeg might promote degradation of free deaminases, whilst the Pepper-bearing gRNA would stabilize these deaminases at editing sites.

Before implementing this design, we speculated that tDeg would need to be sufficiently potent to drive rapid degradation of its fusion partner. Thus, we began by optimizing tDeg using a fluorescent protein-based reporter assay. We fused modified tDegs to the C-terminus of mNeonGreen^40^, using mScarlet3^41^, encoded by the same mRNA, as an internal control (Fig. 1a). A degradation/stabilization index was determined by the ratio of mNeonGreen+ cells to mScarlet3+ cells determined following a flowcytometry assay (Extended Data Fig. 1a-1d).

**Fig. 1.**
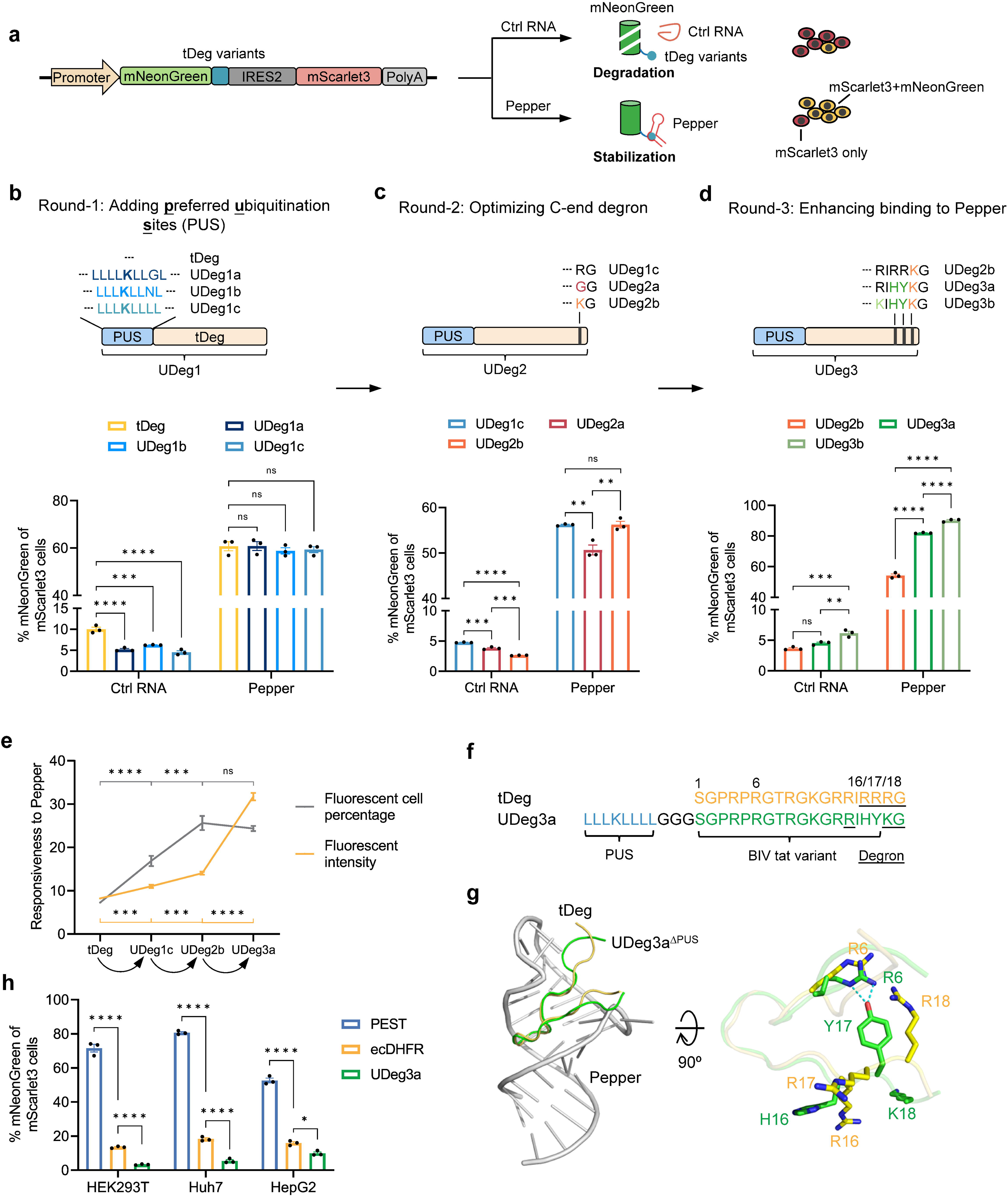
Optimized UDeg-Pepper systems for RNA-regulated protein stability. **a**, Schematic showing assessment of tDeg variants. Fusion of mNeonGreen with tDeg mutants is unstable unless tDegs were bound to their RNA aptamer, Pepper. The same RNA encoded mScarlet3, which was used as an internal control. mScarlet3 single-positive and mScarlet3 mNeonGreen double-positive cells are colored in red and in yellow, respectively. The mNeonGreen/mScarlet3 ratio reflects how stable/unstable the protein is. **b**–**e**, Stepwise optimization of tDeg generates UDeg variants with improved destabilization ability or responsiveness to Pepper. Round-1: Supplement of PUS promotes rapid proteolysis (**b**). Round-2: Replacement with novel C-terminal degrons enhances degradation (**c**). Round-3: Chimeric tat peptides ending with HY heighten the responsiveness to Pepper (**d**). The responsiveness to Pepper is increased at each step (**e**). Responsiveness is defined by the fold change of fluorescent cell percentage or fluorescent intensity in the presence of Pepper. Data are presented as mean ± SEM from 3 biological repeats. One-way ANOVA: ns, not significant, ** *p*<0.01, \*\*\**p*<0.001, **** *p*<0.0001. **f**, UDeg3a is composed of a PUS, a chimeric tat peptide, and an optimized C-terminal degron (underlined residues). The residues are numbered from the beginning of the tat peptide. Tat peptides and degrons involved in binding to Pepper are highlighted in yellow (tDeg) and in green (UDeg3a), respectively. **g**, Structural alignment between tDeg and UDeg3a^ΔPUS^, generated by alphafold3. Y17 forms two hydrogen bonds with R6, potentially stabilizing the UDeg3a conformation to favor its binding to Pepper. The colors and numbers are specified in (F). **h**, UDeg3a is an efficient C-end degron. Data are presented as mean ± SEM from 3 biological repeats. One-way ANOVA: * *p*<0.05, **** *p*<0.0001.

According to the tripartite degron model, protein turnover is regulated by three key components: the E3 ligase recognition motif, an accessible acceptor lysine for ubiquitination, and a disordered degradation initiation site^42^. Beyond the E3 ligase binding motif, the accessibility of the acceptor lysine is crucial for degradation efficiency^42, 43^. Therefore, we hypothesized that adding a ubiquitinable lysine adjacent to tDeg would enhance proteolysis. Previous studies have shown that residues surrounding ubiquitinated lysines are enriched with aromatic and hydrophobic amino acids^42^, especially leucine^44^, and that these ubiquitination sites are typically close to the degron^42^. To enhance degradation, we introduced a stretch of seven leucines centered around a lysine residue immediately upstream of tDeg. We also substituted leucine at the +3 position relative to lysine with either glycine or asparagine (Fig. 1b), both of which were found to be overrepresented in ubiquitination peptides^42^. As expected, all three preferred ubiquitination sites (PUS) significantly promoted mNeonGreen degradation, generating a new group of degrons we termed UDeg. For example, the proportion of mNeonGreen+ in mScarlet3+ cells decreased from 9.97% to 4.52% for tDeg and UDeg1c, respectively. Importantly, UDeg1s were stabilized by Pepper to comparable expression levels (Fig. 1b and Extended Data Fig. 1b). These findings suggest that the PUS facilitated but did not initiate protein degradation, prompting us to select UDeg1c for the next round of optimization.

Recent studies have identified numerous C-terminal degrons via high-throughput screening of the peptidome, revealing that proteins ending in -GG or -RxxxKG motifs were depleted more than other types in the library^45, 46^. We mutated the terminal two -RG residues in UDeg1c to either -GG or -KG to generate UDeg2a and UDeg2b, respectively. Correspondingly, both UDeg2a and UDeg2b exhibited lower basal expression level of mNeonGreen than UDeg1c, with UDeg2b being most effective (Fig. 1c and Extended Data Fig. 1c). Moreover, the mutation in UDeg2b did not affect the stabilizing effect of Pepper. Hence, we selected UDeg2b for subsequent experiments.

In addition to degron optimization, it would also be beneficial to enhance the stabilizing effect of Pepper. We investigated the use of a tat peptide from the Jembrana disease virus (JDV)^47^, previously shown to have a high binding affinity for the bovine immunodeficiency virus (BIV) TAR and its derivatives, such as Pepper^48^. Specifically, the terminal four residues (KIHY) of the tat peptide are responsible for stronger binding to TAR^47^. Based on UDeg2b, we designed two new degrons, UDeg3a and UDeg3b ending in -RIHYKG and -KIHYKG, respectively. Both degrons led to a significantly higher percentage of mNeonGreen+ cells in the presence of Pepper compared to the previous degron, up to 90.3%, indicating much stronger protection by Pepper. Ultimately, we selected UDeg3a over UDeg3b due to its relatively lower basal expression level (Fig. 1d and Extended Data Fig. 1d).

Collectively, we calculated the responsiveness of tDeg, UDeg1c, UDeg2b, and UDeg3a to Pepper in each step, indexed by the fold change in the percentage of mNeonGreen+ cells, which was 7.27±0.19, 16.88±1.18, 25.64±1.66, and 24.39±0.62, respectively. Both UDeg2b and UDeg3a exhibited approximately a 3.5-fold increase compared to tDeg. When based on fluorescent intensity, the responsiveness of these degrons were 8.24±0.1, 11.03±0.37, 14.09±0.33, and 31.76±0.89, respectively (Fig. 1e). UDeg3a, comprised of a PUS, a chimeric tat peptide, and an optimized C-terminal degron (Fig. 1f) had the highest (approx. 3.9 times) increase in fluorescent intensity compared to that of tDeg, making it the prime candidate for selection.

To investigate the enhanced responsiveness of UDeg3a to Pepper, we constructed a structural model of the UDeg3a-Pepper complex using AlphaFold3^49^. The PUS was excluded as it is not involved in Pepper binding. To directly compare interaction patterns, tDeg was superimposed onto UDeg3a. Structural analysis revealed that both UDeg3a and tDeg adopt similar backbone conformations when binding to Pepper, forming a β-hairpin structure that fits into the major groove of Pepper. These results align with the BIV and JDV tat structures previously determined by nuclear magnetic resonance (NMR) spectroscopy^50^, supporting the reliability of the AlphaFold3 model. Notably, the primary structural differences were observed in the peptide termini, where Y17 (numbered from the start of the tat peptide) in UDeg3a formed hydrogen bonds with R6. This intramolecular interaction might help to constrain the conformation of UDeg3a, reducing the free energy of binding to Pepper. Although potential contact between the aromatic ring on Y17 and the RNA base was not explicitly observed in this model (Fig. 1g), such an interaction could further strengthen UDeg3a-Pepper binding^50^. Additionally, electrostatic potential mapping of the peptide surfaces indicated that mutations R16H, R17Y, and R18K reduced the positive charge on tDeg, potentially mitigating electrostatic repulsion between peptide termini during β-hairpin formation (Extended Data Fig. 1e).

We also compared UDeg3a to two commonly used protein degradation domains, PEST^51^ and ecDHFR^52^. We found that UDeg3a was more efficient in promoting proteolysis in all cell types tested (Fig. 1h). The superior destabilization activity and smaller size highlight the usefulness of UDeg3a as a protein degradation tag (Extended Data Fig. 1f). Furthermore, this degradation process appears to be mediated by the ubiquitin-proteasome system (UPS), as UPS inhibition by MG132^53^ resulted in dose-dependent protein stabilization (Extended Data Fig. 1g). Taken together, we have engineered a novel Pepper-regulated protein degradation tag, UDeg3a, enabling both accelerated degradation and enhanced responsiveness to Pepper.

### Development of a conditionally stable ADAR1d for programmable RNA editing

We next sought to utilize Pepper-UDeg3a for generating an RNA base editor with gRNA-regulated stability. As mentioned above, UDeg3a was fused to the C-terminus of the ADAR1 deaminase domain carrying a hyperactive mutation (ADAR1d^E1008Q^, referred to as ADAR1d*)^54^ to induce destabilization. To complement this system, we designed a gRNA incorporating the Pepper domain and a targeting sequence, analogous to the direct repeat and spacer elements of CRISPR gRNAs (Fig. 2a). Crucially, we speculated that Pepper not only recruited but also stabilized ADAR1d*-UDeg3a. As anticipated, we observed destabilization of ADAR1d*-UDeg3a, which can be restored by Pepper. Remarkably, circular gRNA generated via the Tornado system^55^ exhibited stronger stabilizing effects than its linear counterpart (Fig. 2b and Extended Data Fig. 2a), consistent with the increased stability of circular RNAs. As a result, we successfully developed a conditional deaminase system with circular gRNA-regulated stability.

**Fig. 2.**
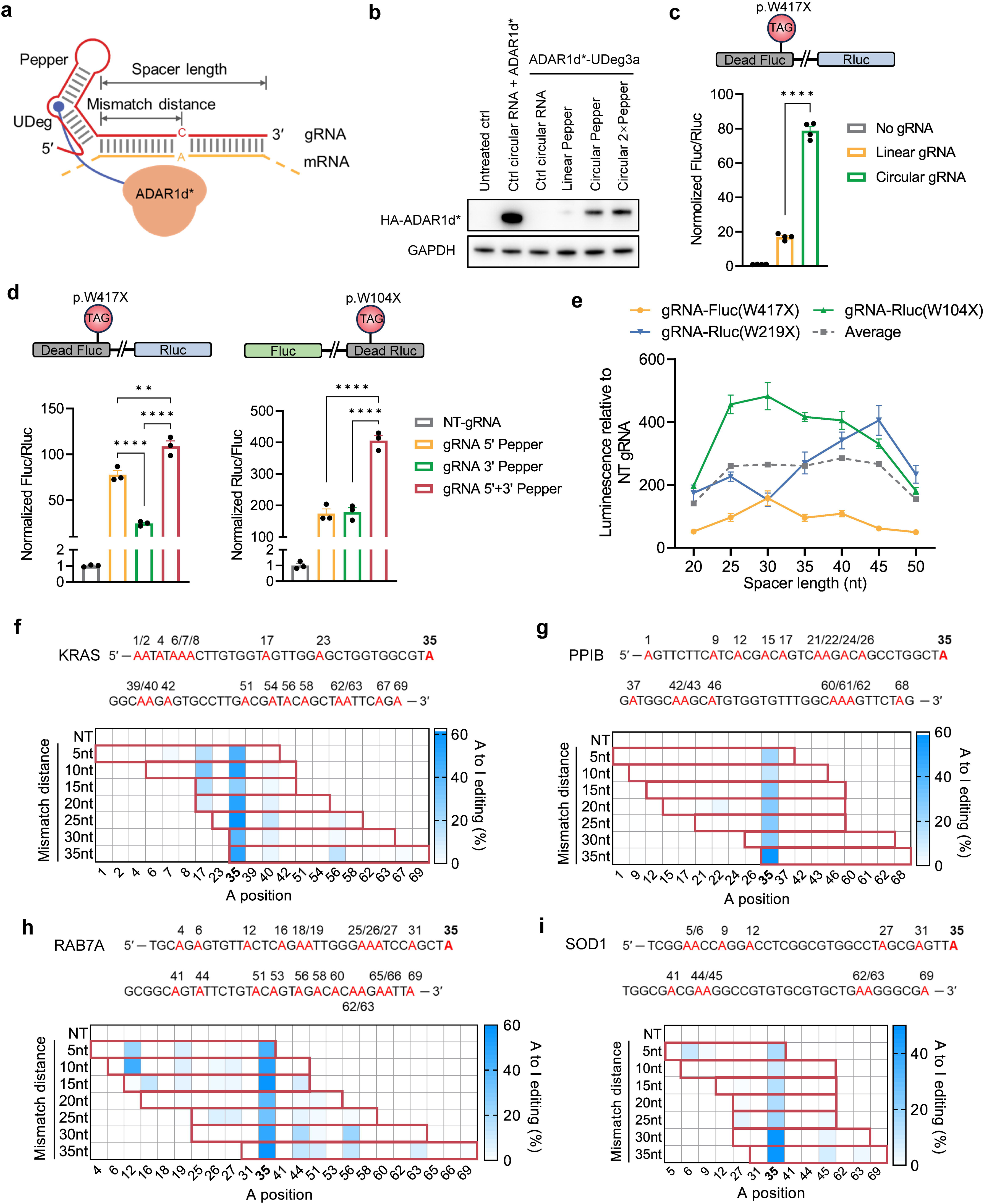
Design and characterization of circular RNA-guided-and-stabilized ADAR1d for RNA editing. **a**, Schematic of an RNA base editor built upon UDeg-Pepper. ADAR1 deaminase^E1008Q^ (referred to as ADAR1d*) fused with UDegs can be directed to target RNA through Pepper-bearing gRNA. **b**, Pepper-regulated stability of ADAR1d* fused with UDeg3a. Pepper within circular RNA is much more efficient than its linear counterpart in stabilizing ADAR1d*-UDeg3a. **c**, Circular gRNA-dependent restoration of the luciferase gene. Data are presented as mean ± SEM from 4 biological repeats. One-way ANOVA: **** *p*<0.0001. **d**, Evaluation of the positional effects of Pepper within gRNA. Data are presented as mean ± SEM from 3 biological repeats. One-way ANOVA: ** *p*<0.01, **** *p*<0.0001. **e**, Optimization of the spacer length. gRNAs with varying spacer lengths were targeted to three sites in the luciferase genes. Data are presented as mean ± SEM from 3 biological repeats. **f**–**i**, Characterization of RECODEv1 composed of ADAR1d*-UDeg3a and circular gRNA with a 40-nt spacer flanked by bilateral Peppers. Heatmaps show the editing rates of all adenosines, highlighted in red, covered by guides with variable mismatch distances targeting endogenous transcripts of *KRAS*(**f**), *PPIB*(**g**), *RAB7A*(**h**), and *SOD1*(**i**). The target adenosines are emphasized in bold. For each guide, the covered adenosines are outlined in red in the heatmaps. Angled numbers below heatmaps indicate nucleotide positions from 1 to 69. Data are presented as mean ± SEM from 3∼4 biological repeats.

To evaluate RNA editing activity, we used this system to correct nonsense mutations (TGG to TAG or TGA codons) in luciferase genes via A-to-I editing. The mutation sites were specified in each experiment, with another intact luciferase gene serving as the internal control (Extended Data Fig. 2b). Our initial design featured a 30-nt spacer with a central cytidine pairing to the target adenosine. Following co-transfection into HEK293T cells, we observed gRNA- and ADAR1d*-UDeg3a-dependent restoration of firefly luciferase (Fluc) activity (Fig. 2c and Extended Data Fig. 2c). Notably, circular gRNA was significantly more efficient than its linear counterpart, likely due to the increased abundance of both gRNA and ADAR1d*-UDeg3a (Fig. 2b). Thus, circular gRNA was employed in subsequent experiments.

In accordance with the enhanced binding between UDeg3a and Pepper, ADAR1d*-UDeg3a exhibited greater activity than ADAR1d*-UDeg2b in restoring the mutated *Fluc* gene. Moreover, ADAR1d* outperformed ADAR2d* (ADAR2 deaminase domain with a E488Q mutation) ^56^, implying that ADAR2d* may be less compatible with the UDeg-Pepper system in this context (Extended Data Fig. 2c). We next explored the influence of subcellular localization of deaminases on editing activity. Luciferase assays revealed that ADAR1d*-UDeg3a lacking localization signals performed better than variants with a nuclear localization signal (NLS) or a nuclear export signal (NES) (Extended Data Fig. 2c). Similar trends were observed during the editing of two endogenous genes, *ALDOA* and *DAXX* (Extended Data Fig. 2d).

After establishing the protein component of the RNA editor, we turned to optimizing the gRNA design. We investigated the positional effects of Pepper within gRNAs targeting different reporters (*Fluc* or Renilla luciferase [*Rluc*] genes). We found that placing Pepper at both ends of the spacer led to greater editing efficiency than positioning it solely at the 5′ or 3′ end. This aligns with previous reports that increasing recruitment elements in gRNA often improves editing efficiency^33, 57, 58^, potentially amplified in our system by the resulting stabilization of deaminases. Among RECODE gRNAs with a single Pepper, a 5′ terminal position was preferred, at least in targeting Fluc (Fig. 2d).

To further refine the system, we examined the optimal spacer length using bi-Pepper gRNAs with spacers ranging from 20 to 50 nt, in 5-nt increments. Luciferase assays indicated that spacer lengths between 25 and 45 nt offered reliable editing, with the averaged activity of three different test gRNAs peaking at 40 nt (Fig. 2e). Based on these findings, we adopted a gRNA design featuring a 40-nt spacer flanked by Pepper domains at both ends. This optimized gRNA architecture, combined with ADAR1d*-UDeg3a, established a new platform: RECODE version 1 (RECODEv1).

### Characterization of RECODEv1

To further evaluate RECODEv1, we examined the impact of A−C mismatch position on editing efficiency. The mismatch position was defined as the distance from the 5′ end of the spacer. We systematically tiled gRNAs across target adenosines in both endogenous (*KRAS*, *PPIB*, *RAB7A*, *SOD1*) and transfected (*Angptl3*, *Idua*) transcripts, with a 5-nt step, in HEK293T cells. All adenosines within the gRNA footprint were analyzed to comprehensively map the editing preferences of RECODEv1. We found that target adenosines were reliably edited by all gRNAs, regardless of mismatch position, though the efficiency varied (Fig. 2f-2i, Extended Data Fig. 2e-2g).

For most sites (*PPIB*, *RAB7A*, *SOD1*, and *Angptl3* site 1), the mismatch position at 35 nt yielded the highest editing efficiencies, ranging from 49% to 62%. However, this trend was not universal. For *Angptl3* site 2 and *Idua*, the mismatch position of 20 nt appeared to be optimal (Extended Data Fig. 2h). Based on these observations, we recommend starting mismatch positions of 20 or 35 nt for new applications, with the caveat that site-specific optimization may be necessary.

Additionally, the editing profiles revealed bystander editing at the target loci that was confined to regions covered by the gRNA spacers (Fig. 2f-2i, Extended Data Fig. 2e-2g). This corresponds with the established preference of ADAR deaminases for adenosines within double-stranded RNA (dsRNA) regions^59^. Therefore, the shorter spacers required by RECODEv1 inherently promote more precise editing compared with editing systems needing much longer gRNAs. Moreover, bystander editing could potentially be minimized by introducing A−G mismatches^18^, single-adenosine bulges^19^, or G−U wobble base pairs^23^.

### Development of RECODEv2 for enhanced specificity

The development of RECODEv1 established a system where the stability of ADAR1 deaminase could be regulated by gRNAs. To further improve the performance of this approach, we aimed to design a gRNA with target RNA-activated Pepper. Ideally, deaminases would be stabilized only at editing sites. Drawing inspiration from molecular beacons^60^, which are RNA imaging tools that transition from a “closed” to an “open” state upon hybridization with their targets, we introduced a sequence complementary to the 3′ end of the spacer downstream of Pepper, forming a stem-loop structure. Extending the stem to Pepper may interfere with its folding^61^. However, the presence of target RNA would open the inhibitory stem by forming a spacer-target duplex, thereby restoring Pepper’s active conformation (Fig. 3a).

**Fig. 3.**
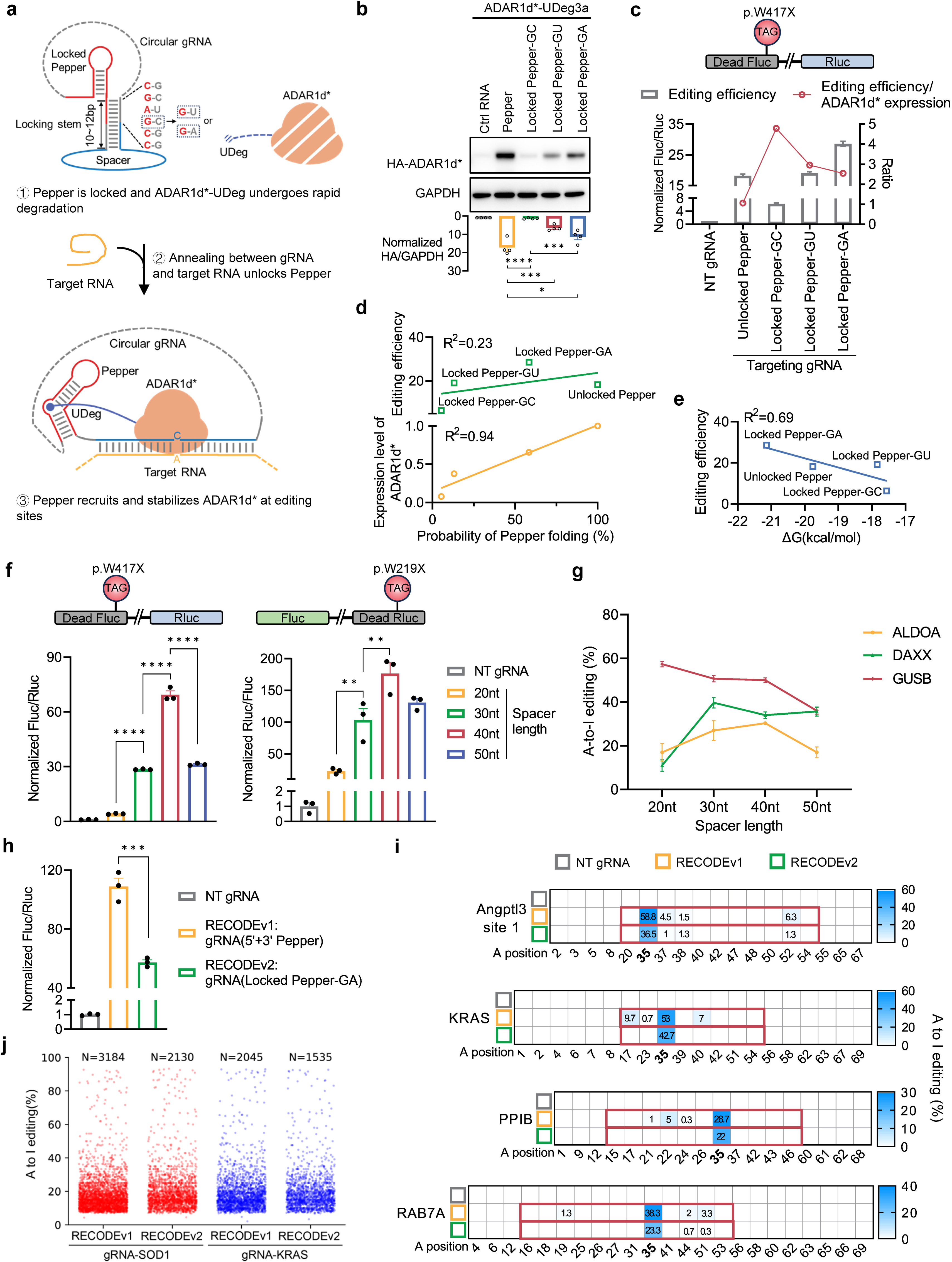
Target-switchable gRNA for enhanced specificity. **a**, Schematic showing switchable-by-target strategy for improving RECODE specificity. IZ Pepper is locked by an inhibitory stem, which prevents the stabilization of ADAR1d*-UDeg3a. IZ Hybridization between guides and target RNA unlocks Pepper. IZ ADAR1d*-UDeg3a is recruited and stabilized at editing sites. For optimal usage, the strength of the inhibitory stem should be appropriately determined. **b**, Expression levels of ADAR1d*-UDeg3a are affected by the inhibitory stem. Locking Pepper with a fully base-paired stem almost abolished the gRNA’s stabilizing effect. Weakening the inhibitory stem gradually increases ADAR1d*-UDeg3a expression. The expression levels of ADAR1d*-UDeg3a are represented by the band intensity in Western blot, which are normalized to the group of ctrl RNA. **c**, Differences in inhibitory stems can markedly affect RNA editing activity and the ratio of RNA editing activity to ADAR1d* expression level. RNA editing efficiency is represented by the normalized bioluminescence, which is presented as mean ± SEM from 4 biological repeats. The ratio is calculated as bioluminescence value divided by ADAR1d*-UDeg3a expression level in (B). **d**,**e**, Two thermodynamic parameters of gRNA correlated with ADAR1d*-UDeg3a expression level and editing efficiency. The probability of Pepper folding (**d**) was strongly correlated with ADAR1d*-UDeg3a expression levels (normalized to unlocked Pepper), but not with editing efficiency. Linear regression analysis (**e**) revealed a moderate correlation between guide-target binding free energy and editing efficiency. **f**, Spacer length optimization of gRNA with locked Pepper-GA. **g**, Editing of endogenous transcripts confirmed an optimal spacer length of approximately 30∼40nt. **h**, Comparison between luciferase assays of RECODEv1 and RECODEv2 gRNAs. **i**, Heatmaps showing the editing rates of adenosines within a window of 69 nt centered on target adenosines. These rates were assessed following treatment with RECODEv1 and RECODEv2, with spacer-matched gRNAs. Adenosines within a guide-target duplex are outlined in red. The angled numbers indicate nucleotide positions from 1 to 69. Values are shown as means from 3 biological repeats. **j**, Transcriptome-wide analysis of off-target effects of RECODEv1 and RECODEv2 with spacer-matched gRNAs. The number of global A-to-I edits is shown for each condition. For **b**, **f**, and **h**, data are presented as mean ± SEM from 3∼4 biological repeats. One-way ANOVA: * *p*<0.05, ** *p*<0.01, *** *p*<0.001, **** *p*<0.0001.

We positioned the switchable Pepper at the 5′ end of the spacer, as it demonstrated higher activity in this configuration (Fig. 2d). The inhibitory stem was designed to be 10∼12 bp long, with a 6-bp upper segment directly competing with Pepper folding. We reasoned that the strength of this inhibitory stem, or “Pepper lock”, would be critical for efficient transitions between the active and inactive states. Thus, we generated several gRNAs with varying stem strength, via introduction of a G−U wobble base pair or G−A mismatch (Fig. 3a). We evaluated the impact of stem strength on ADAR1d*-UDeg3a stability and editing efficiency, with a 30-nt spacer targeting Fluc. The results indicated that a fully base-paired stem severely restrained the protein levels of ADAR1d*-UDeg3a, suggesting a tightly locked Pepper (Fig. 3b and Extended Data Fig. 3a). Gradual weakening of the locking stem increased both ADAR1d*-UDeg3a levels and editing efficiency. Notably, the ratios of editing activity to ADAR1d*-UDeg3a expression levels increased across all three gRNAs with locked Pepper, implying improved specificity. Interestingly, locked Pepper with G−A mismatch resulted in even higher editing efficiency than the unlocked Pepper, despite lower expression levels of ADAR1d*-UDeg3a (Fig. 3b and 3c). This construct was therefore selected as it allowed for the minimization of deaminase expression levels without compromising on-target activity. However, if specificity is a priority, the other two gRNAs with locked Pepper are viable alternatives.

To obtain deeper understanding of the relationship between gRNA folding, ADAR1d*-UDeg3a expression, and editing efficiency, we sought to identify correlative thermodynamic parameters. We excluded the minimum free energy (MFE) of the locking stem, as it did not account for loop structures. Instead, we predicted the folding of the entire gRNA sequence (excluding circularization elements) and calculated the partition function using RNAfold program^62^. Our analysis revealed a strong correlation (R²=0.94) between the probability of Pepper folding—defined by the pairing likelihood of the terminal base pair in Pepper—and the expression levels of ADAR1d*-UDeg3a, underscoring the viability of precisely modulating ADAR1d*-UDeg3a expression through gRNA engineering. In contrast, the correlation between Pepper folding probability with editing efficiency was notably weaker (R² = 0.23), tapering off before the peak of ADAR1d*-UDeg3a expression was reached (Fig. 3d and Extended Data Fig. 3b). This discrepancy also supports the scenario of efficient transition of Pepper from the locked to the active state upon hybridization of the guide with the target.

The weak correlation between Pepper folding dynamics and editing efficiency suggests the necessity of considering target RNA. To explore this, we calculated the free energy of guide-target binding as a measure of hybrid stability (Extended Data Fig. 4). The analysis revealed that subtle modifications to the stem can affect the energy required for loop annealing to the target. Remarkably, there was a moderate correlation between binding free energy and editing efficiency (R² = 0.69) (Fig. 3e). These findings further emphasize the critical role of fine-tuning stem strength and its predictable impact on editing yields.

Next, we optimized spacer length using locked Pepper with a G−A mismatch. Luciferase assays revealed that a 40-nt spacer was optimal for both *Fluc* and *Rluc* gRNAs, a result corroborated by editing endogenous transcripts such as *ALDOA*, and *DAXX* (Fig. 3f and 3g). Peak editing efficiency generally occurred with spacers between approximately 30−40 nt, though an exception was observed where a 20-nt spacer maximized editing for *GUSB* (Fig. 3g).

Previous studies show that addition of a flexible RNA linker into circular gRNA contributes to release of the conformation constrains and boosts editing efficiency.^19, 25^ Hence, the gRNA used above carried a flexible AC linker downstream of the spacer. Next, we tried to append AC linkers to both the 5′ end of Pepper and the 3′ end of the spacer and found this modification further enhanced editing activity (Extended Data Fig. 5a, and 5b). Collectively, the gRNA architecture comprising locked Pepper with a G−A mismatch, a 40-nt spacer, and AC linkers—was designated RECODEv2.

### Comparison between RECODEv1 and RECODEv2

We compared RECODEv2 to RECODEv1 in terms of editing efficiency, bystander editing, and global off-target effects using spacer-matched gRNAs. The luciferase assays indicate a significant reduction of activity in RECODEv2 compared to RECODEv1 (Fig. 3h). Analysis of editing rates also indicated lower activity of RECODEv2 in targeting Angptl3, KRAS, PPIB, and RAB7A transcripts (Fig. 3i), including bystander editing, which was almost eliminated in the cases of KRAS and PPIB. This corresponds with our results in which Peppers flanking the spacer at both ends induced more edits than single Pepper (Fig. 2d). Transcriptome analysis further revealed fewer off-target edits (Fig. 3j), which may result from the attenuated effect of RECODEv2 gRNA in stabilizing ADAR1d*-UDeg3a (Fig. 2b and 3b).

### Engineering ADAR1d for enhanced activity

Developing highly active deaminases is crucial for targeting editing-resistant sites. Alongside efforts to confer conditional stability on ADAR1d*, we sought to further enhance its catalytic activity through protein engineering. Given the absence of structural information for ADAR1d, we utilized AlphaFold3^49^ to model the ADAR1d*-RNA duplex, identifying several regions that interact directly with the RNA substrate. One region of particular interest was a loop spanning residues 972 to 999, homologous to the 5′ binding loop of ADAR2 deaminase^26, 63^. This loop has been proposed as the structural basis for the differing substrate specificities between ADAR1 and ADAR2. Moreover, its primary sequence is highly variable among ADAR1 orthologs across diverse species, including fish, reptiles, and birds (Fig. 4a). Considering the broad impact of temperature on both enzymatic catalysis and dsRNA formation, we hypothesized that the ADAR1 deaminase domain may have evolved distinct functional characteristics in poikilotherms and birds, whose body temperatures differ significantly from those of mammals. This prompted us to substitute the loop in ADAR1d* with orthologous counterparts from these species to identify hyperactive variants.

**Fig. 4.**
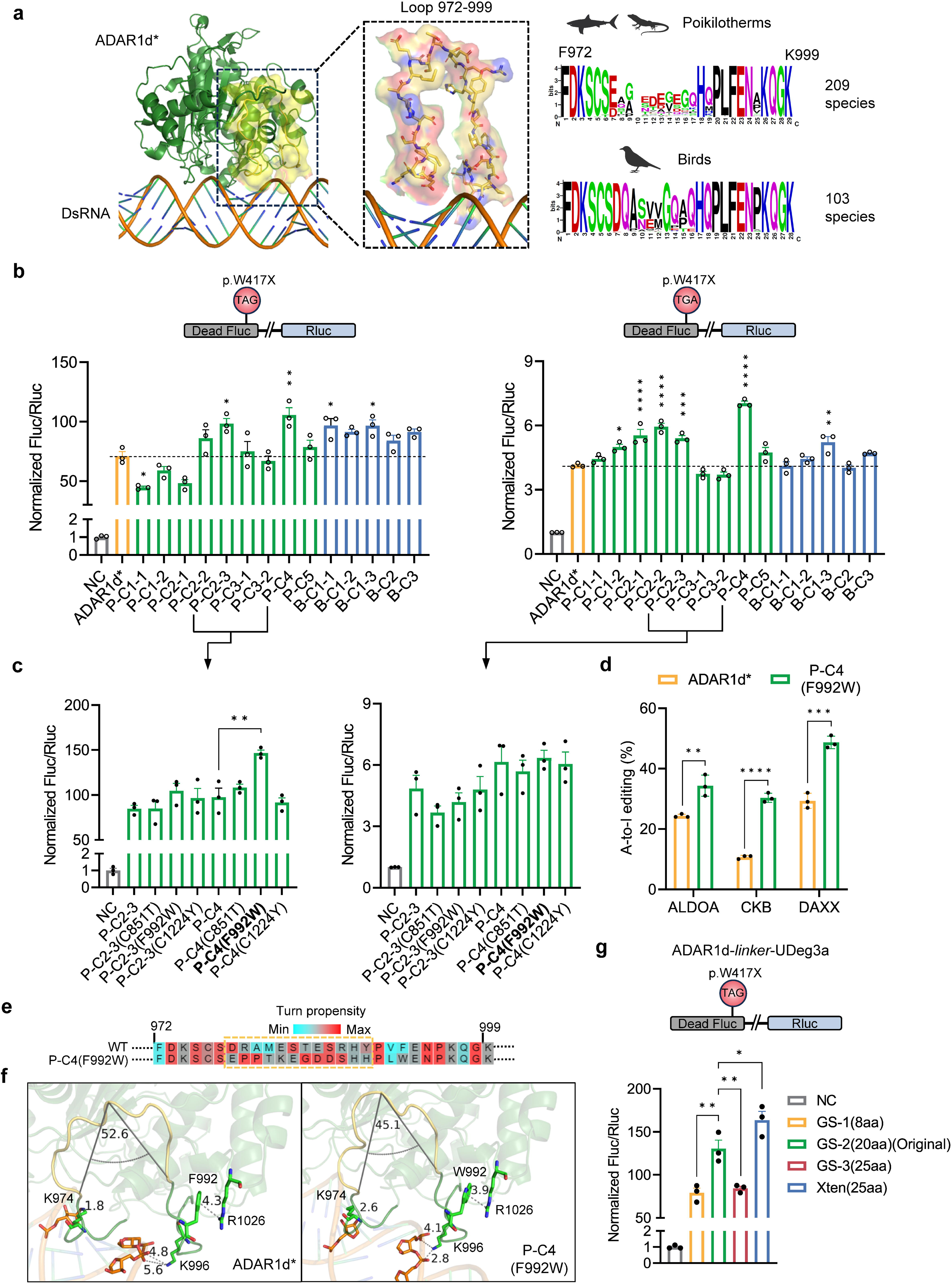
Engineering ADAR1d for increased activity through structure-guided consensus. **a**, Illustration of the 5′ binding loop of ADAR1d* in a structure model by Alphafold3 and its variation among orthologs. The termini of this loop made direct contact with dsRNA substrate, as shown by the loop surface in yellow. **b**, Evaluation of ADAR1d mutant activity on editing either UAG (left) or UGA (right) in luciferase assays. The mutants were generated by exchanging the 5′ binding loop with consensus sequences of those from poikilotherm vertebrates and Aves. P-C2-3 and P-C4, which carried loops from fish, were most active and were selected for subsequent applications. NC represents negative control of no deaminase. **c**, Identification of a hyperactive ADAR1d mutant through rational mutagenesis. Mutation of F992W conferred extra activity on P-C4 in the editing of UAG, which is highlighted in bold. **d**, Editing of endogenous transcripts confirmed that P-C4 (F992W) had higher activity than ADAR1d*. **e**, Pairwise alignment of the 5′ binding loop from WT ADAR1d and P-C4 (F992W); Red color represent high turn propensity and cyan colors represent low turn propensity. Highly variable residues among species are boxed in yellow. **f**, Structural analysis of the increased activity of P-C4 (F992W). The structures of ADAR1d* and P-C4 (F992W) were generated by Alphafold3, with a close-up view of 5′ binding loop. The turn in yellow corresponds to boxed residues in (E). **g**, Linker optimization between P-C4 (F992W) and UDeg3a. For **b**, **c**, **d**, and **g**, data are presented as mean ± SEM from 3 biological repeats. One-way ANOVA: * *p*<0.05, ** *p*<0.01, *** *p*<0.001, **** *p*<0.0001.

To streamline the screening process, we generated consensus sequences for the ADAR1 5′ binding loop from 209 poikilotherms and 103 bird species, in which the residue frequencies at each position might reflect evolutionary fitness (Extended Data Fig. 6). The sequences from each species can be classified into eight clusters based on the similarity. From these clusters, we derived eight consensus motifs: P-C1∼5 from poikilotherms and B-C1∼3 from birds. We then selected 1–3 representative sequences from each consensus motif and replaced the native loop in ADAR1d* with these sequences (Extended Data Fig. 7a). The activity of the resulting variants was evaluated by testing their ability to restore two termination codons (UAG and UGA) in *Fluc*. Most variants exhibited activity comparable to the original ADAR1d*-UDeg3a, and only one substitution slightly impaired UAG editing, indicating the overall functional fitness of nature-derived sequences. Notably, two variants, P-C2-3 and P-C4, demonstrated significantly enhanced activity in editing both UAG and UGA and were selected for further analysis (Fig. 4b).

Inspired by prior high-throughput mutagenesis studies^26, 64^, we introduced three additional mutations into P-C2-3 and P-C4. Among these, the F992W mutation notably increased the activity of P-C4 in UAG editing. However, this enhancement was not observed for UGA, likely due to a smaller dynamic range for this codon (Fig. 4c). We then compared the performance of P-C4 (F992W) with ADAR1d* in editing endogenous transcripts, with gRNAs of RECODEv2. The P-C4 (F992W) variant consistently outperformed ADAR1d*, achieving up to a threefold increase in editing rates across three tested sites (Fig. 4d).

To gain structural insights into these hyperactive mutants, we aligned the mutated sequence with the wild type and generated structural models using AlphaFold3^49^. The analysis revealed that the central segment of the mutated loop was enriched in residues with high turning propensity (Figure 4e). Specifically, four amino acids (K982, E983, G984, D985) formed a classical β-turn at the loop apex, facilitated by electrostatic interactions between K982 and D985 (Extended Data Fig. 7b). This structural modification reduced the loop angle, positioning K996 closer to the phosphate backbone of the substrate RNA. Additionally, a cation-π interaction was observed between F992 and R1016 in ADAR1d*. The F992W mutation appeared to strengthen this interaction because of the larger π-electron cloud of tryptophan, thereby further stabilizing the loop structure (Fig. 4f).

Finally, we optimized the linker between P-C4 (F992W) and UDeg3a, identifying the xten linker as the most effective in promoting RNA editing (Fig. 4g, and Extended Data Fig. 7c). Consequently, P-C4 (F992W) and the xten linker were integrated into the RECODE platform for subsequent applications.

### RECODE compares favorably to other established technologies for RNA editing

We then benchmarked RECODE against other well-established RNA editing platforms including REPAIRv2 (CRISPR/Cas13-based RNA editing)^34^, MCP-ADAR1d*^33^, LEAPER2.0^19^ or cadRNAs^18^ (Circular gRNAs recruiting endogenous ADARs). To facilitate a fair comparison, each method was implemented using its optimal gRNA architecture. For REPAIRv2, the gRNA was designed with a 50-nt spacer and a 35-nt interval between mismatched cytosine and direct repeats. In the case of MCP-ADAR1d*, the gRNA consisted of a 22-nt spacer flanked by two MS2 structures, with an opposite cytosine positioned 7 nt from the 5’ MS2 element. LEAPER2.0 utilized a 151-nucleotide target sequence containing a centrally positioned opposite cytosine flanked by AC linkers. With regard to RECODEv2, a 40-nt spacer with central opposite cytosine was used.

To assess both on-target and bystander editing, we quantified the editing rates of the target adenosine in addition to surrounding adenosines within a 151-nucleotide window centered on the target, across a range of transfected and endogenous transcripts. We found consistently high editing efficiencies across these sites with MCP-ADAR1d* and RECODEv2. In contrast, there was a marked dependence on the specific editing sites in the editing activity of REPAIRv2 and LEAPER2.0 (Fig. 5a-5f). Notably, RECODEv2 achieved robust editing at DAXX and Rluc (W104X), with editing rates of 71% and 82.3%, respectively, whereas editing at these loci was undetectable for REPAIRv2 and LEAPER2.0 (Fig. 5a and 5e). Furthermore, there were markedly fewer bystander effects during RECODEv2 editing compared to MCP-ADAR1d* (Figures 5A-5F), which is also shown by the ratio of on-target to cumulative bystander (i.e., the sum of all bystander editing events within the window) editing rates (Fig. 5g). Additionally, we plotted on-target editing activity against the cumulative bystander editing rates. This analysis revealed that the data points for RECODEv2 predominantly clustered in the upper-right quadrant, again indicating a superior ratio of on-target to bystander editing over the other systems (Extended Data Fig. 8a).

**Fig. 5.**
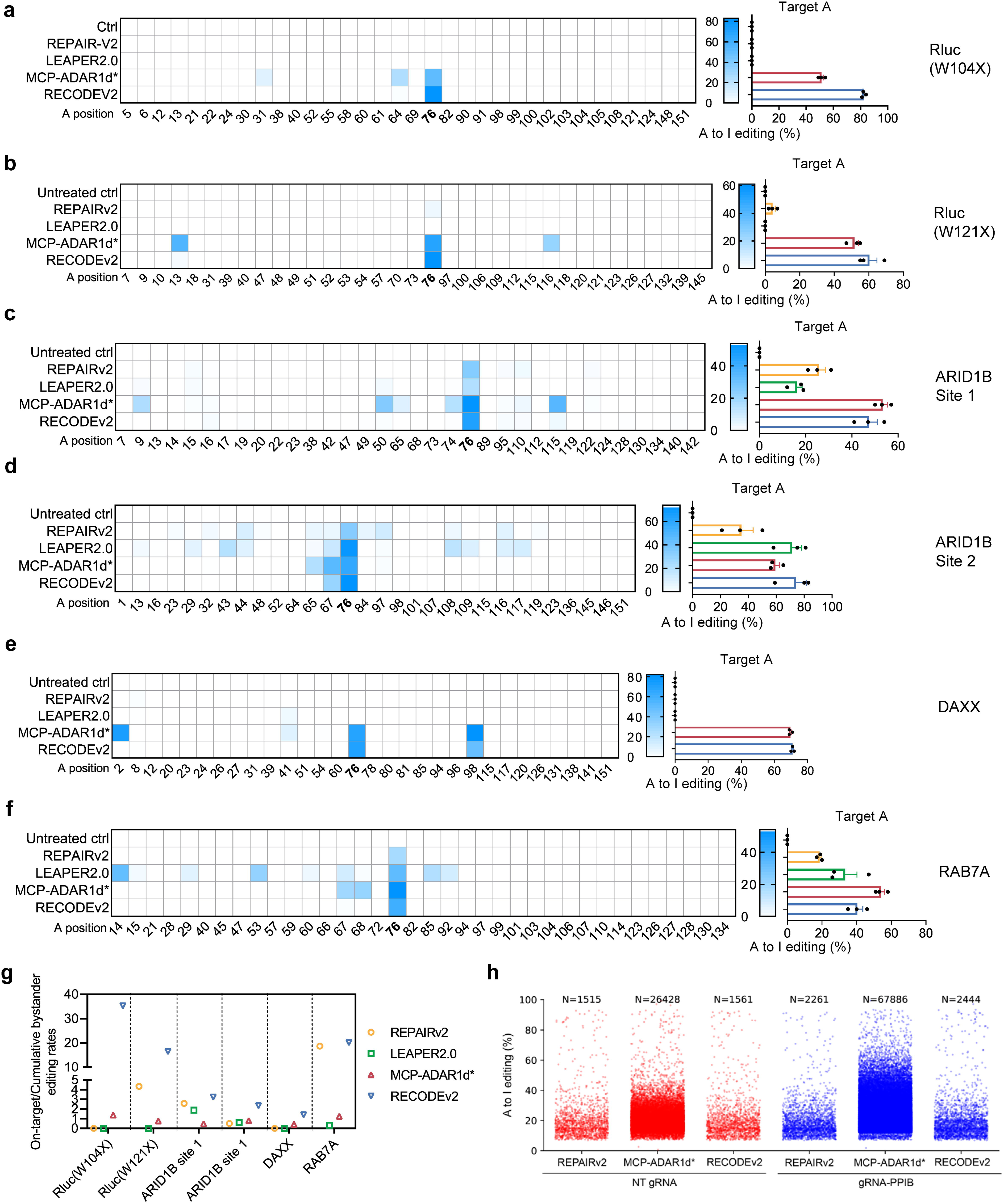
Benchmark RECODE against established RNA editing platforms. **a**–**f**, Heatmaps (left) illustrating the editing rates of adenosines within a 151-nt window centered on the target adenosines in both transfected transcript Rluc (**a**, **f**) and endogenous transcripts ARID1B (**c**, **d**), DAXX (**e**), and RAB7A (**f**). These rates were assessed following treatment with REPAIRv2, LEAPER2.0, MCP-ADAR1d*, and RECODEv2. The angled numbers below each heatmap indicate nucleotide positions from 1 to 151. The target adenosine is numbered 76. Values are shown as means from 3 biological repeats. Histograms (right) illustrating the editing rates of target adenosines. Data are presented as mean ± SEM from 3 biological repeats. **g**, The ratio of on-target to cumulative bystander editing rates is shown for each site. Cumulative bystander editing rates refer to the sum of all bystander editing within the 151-nt windows, which was added by a minimum value to ensure the denominator does not equal zero. **h**, Transcriptome-wide analysis of off-target effects. The number of global A-to-I edits is shown for each condition.

We then conducted transcriptome-wide sequencing to assess the global off-target effects of three exogenous deaminase-dependent methods: REPAIRv2, MCP-ADAR1d*, and RECODEv2. A relatively higher amount (200 ng) of plasmids were used here to emphasize the off-target effect. Our analysis revealed that REPAIRv2 and RECODEv2 induced comparable off-target editing events, both of which were dramatically fewer than those observed with MCP-ADAR1d*. These results are in agreement with previous studies demonstrating that ADAR1d generally produces more extensive off-target edits compared to ADAR2d^33^, with this difference being further magnified by the high-fidelity variant of ADAR2d used here in REPAIRv2^34^. Notably, RECODEv2 substantially mitigated the off-target effects associated with the hyperactive ADAR1d variant, underscoring the efficacy of the conditional stabilization strategy (Fig. 5h). Additionally, motif enrichment analysis of the off-target editing sites revealed a consistent sequence preference, with 5′ W and 3′ S motifs being enriched (Extended Data Fig. 8b).

Beyond the favorable on-target to off-target editing ratio, RECODE possesses two other advantageous features. First, the size of its protein component, ADAR1d*-UDeg3a, is smaller (486aa) than that of MCP-ADAR1d* (587aa) and dPspCas13b-ADAR2^E488Q/T375G^ (1490aa), making it suitable for viral delivery with limited payload capacity. Second, the relatively low immunogenicity, as assessed by the immunogenicity score^65^, suggests a reduced risk of eliciting an immune response (Extended Data Fig. 8c), further enhancing its therapeutic potential.

### Efficient editing of an ALS-relevant *FUS* mutation *in vitro* and *in vivo*

To showcase its therapeutic potential, we used RECODE to target an ALS-relevant mutation in the *FUS* gene. In ALS, a cluster of non-synonymous mutations disrupts the C-terminal NLS of FUS, resulting in cytoplasmic mislocalization^66, 67^. This mislocalization impairs intra-axonal protein synthesis, leading to synaptic dysfunction in neurons^68^. For instance, the c.1555C>T (p.Q519X) mutation truncates the NLS (residues 510–526) ^67^, potentially contributing to cytoplasmic toxicity. Thus, we sought to leverage RECODEv2 to restore this truncated NLS and correct cytoplasmic mislocalization by converting premature termination codons (TAG) to tryptophan codons (TGG).

We first generated SYFP2-tagged FUS (Q519X) to visualize FUS subcellular localization (Figure 6a). Co-delivery of FUS (Q519X) and RECODEv2 to HEK293T cells led to a conversion rate of 90.3% at the target site (Fig. 6b). In accordance with this strikingly high editing yield, we observed a robust rescue of mislocated FUS. The nuclear localization index of FUS, measured by Pearson’s correlation between SYFP2-FUS (Q519X) and DAPI, increased from –0.02 to 0.34, approximating to 0.36 in WT FUS (Figures 6c and 6d). To rule out the influence of SYFP2 on FUS (Q519X) transport, we replaced SYFP2 with a small HA tag, which enabled discrimination from endogenous FUS. After co-transfection of HEK293T cells, nuclear and cytoplasmic proteins were prepared separately. Western blot analysis of cytosolic fractions revealed a significant reduction of HA-FUS with targeting gRNA (Fig. 6e and 6f). Collectively, these cell model assays all accord with the robust activity RECODEv2 in restoring the mislocalization of FUS (Q519X).

**Fig. 6.**
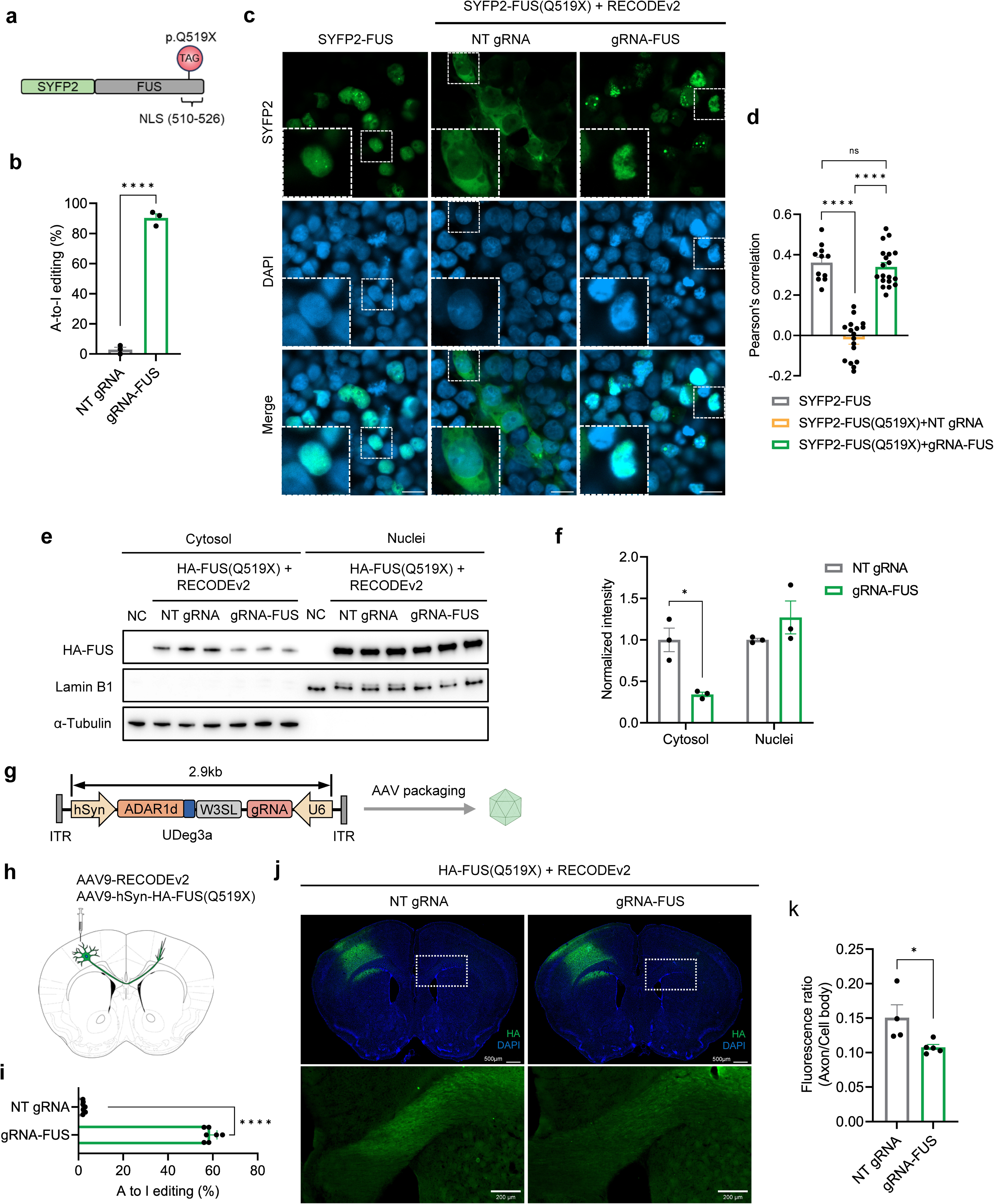
Robust editing of ALS-relevant *FUS* mutation *in vitro* and *in vivo*. **a**, Schematic of FUS mutant (Q519X) fusion to SYFP2. The mutation truncates C-terminal NLS. **b**, Histogram showing the editing efficiency of SYFP2-FUS(Q519X) by RECODEv2 in HEK293T cells. Data are presented as mean ± SEM from 3 biological repeats. Two-tailed *t*-test: **** *p*<0.0001. **c**, Representative images illustrating the subcellular distribution of SYFP2-FUS in HEK293T cells. The nuclei were visualized by DAPI staining. Boxed cells within the white dotted lines are magnified. Scale bar, 20 μm. **d**, Quantitative analysis of nuclear localization of SYFP2-FUS in (C) indicates RECODEv1 rescued SYFP2-FUS(Q519X) mislocalization to the cytoplasm. The nuclear localization is indexed by Pearson’s correlation between SYFP2 and DAPI. Data are presented as mean ± SEM from n=11 for SYFP2-FUS, n=17 for SYFP2-FUS (Q519X) with NT gRNA, n=19 for SYFP2-FUS(Q519X) with targeting gRNA. One-way ANOVA: ns, not significant, **** *p*<0.0001. **e**, Western blot analysis of HA-FUS(Q519X) in cytosolic and nuclear fractions of HEK293T cells. HA-FUS was detected by HA immunoblots. The purity of fractions was determined by immunoblots of Lamin B1 (nuclei) and α-Tubulin (cytosol). **f**, Quantitative analysis of immunoblot bands in (E) reveals RECODEv2 attenuated HA-FUS(Q519X) mislocalization to the cytoplasm. Data are presented as mean ± SEM from 3 repeats. Two-tailed unpaired *t*-test: * *p*<0.05. **g**, Schematic showing RECODE loaded into rAAVs. The total size of RECODE components including cis-regulatory elements is 2.9 kb, far below the rAAV packaging limit. **h**, Schematic illustrating stereotaxic viral injection into the M1. A cluster of M1 neurons send their axons to the contralateral cortex. **i**, Histogram showing the editing efficiency of HA-FUS(Q519X) by RECODEv2 in the mouse brain. Data are presented as mean ± SEM from n=7 animals for each group. Two-tailed unpaired t-test: **** *p*<0.0001. **j**, Representative immunostaining images against HA-FUS(Q519X) showing the whole brain section (top). White dotted boxes indicate signals in axons selected for magnification (bottom). Scale bars, 500 μm (top) and 200 μm (bottom). **k**, Quantitative analysis of images in (J) showed RECODEv2 reduces HA-FUS(Q519X) aggregation in axons, as measured by the ratio of fluorescent intensity in axons to that in cell body. Data are presented as mean ± SEM from n=4 animals for NT, n=5 animals for targeting gRNA. Two-tailed unpaired *t*-test: * *p*<0.05.

Next, we evaluated whether RECODEv2 can facilitate effective rescue *in vivo*. RECODEv2 was packaged into recombinant adeno-associated viruses (rAAVs) that are widely used in gene therapy. The total size of RECODEv2 components, including cis-regulatory elements, was 2.9 kb—well below the 5 kb packaging limit of rAAVs (Fig. 6g) ^69^. We separately packaged RECODEv2 and HA-FUS (Q519X) into AAV9 vectors and co-injected them into the primary motor cortex of mice (Fig. 6h), a region frequently affected in ALS^70^. As a control, RECODEv2 with a non-targeting gRNA was used. Four weeks post-injection, mouse brains were harvested for RNA extraction and tissue sectioning. RECODEv2 with targeting gRNA achieved an editing rate of 58.8% (Fig. 6i). Immunostaining of brain slices revealed significantly less HA-FUS signals in neuronal axons projecting to the contralateral cortex in the RECODEv2 group, as indicated by lower axon-to-cell-body fluorescence intensity ratios (Fig. 6j and 6k). These findings underscore the therapeutic potential of RECODEv2.

### RECODE is broadly applicable to disease-relevant mutations

To demonstrate the broad applicability of RECODE to clinically relevant loci, we applied it to correct multiple mutations in SOD1, another gene implicated in ALS^71^. Seven G-to-A mutations (c.112, c.124, c.280, c.301, c.304, c.335, and c.404) were selected from the ClinVar database^72^ due to their high risk of causing ALS. After co-delivering constructs encoding SOD1 mutants and RECODEv2 into HEK293T cells, we assessed the conversion rates. Efficient editing was observed across these sites, ranging from 45.7% to 91.7%. Notably, six of the seven sites exhibited editing rates exceeding 70% (Extended Data Fig. 9), highlighting the robustness of RECODEv2 in targeting diverse loci.

### Installation of a protective mutation in *Angptl3*

Beyond correcting deleterious mutations, we aimed to use RECODE to install protective mutations in wild-type transcripts. For this, we used ANGPTL3, a hepatically secreted protein that inhibits the clearance of triglyceride-rich lipoproteins and cholesterol in the circulatory system^73, 74^. Previous studies have shown that loss-of-function mutations, such as c.1250A>G (p.Y417C) in *ANGPTL3* are associated with reduced plasma triglyceride levels and a lower risk of coronary artery disease^74^. We used RECODEv1 to introduce this mutation into mouse Angptl3 mRNA. After co-delivering RECODEv1 and HA-tagged Angptl3 into HEK293T cells, we quantified the editing efficiency and analyzed extracellular secretion (Extended Data Fig. 10a). RECODEv1 achieved a high editing efficiency of 64.3% and dramatically reduced extracellular secretion of Angptl3 (an 81.4% reduction) (Extended Data Fig. 10b-10d). These findings demonstrate the potential of RECODE to introduce therapeutic mutations for treating atherosclerotic cardiovascular disease.

### Translational interference via start codon targeting

To expand RECODE’s utility, we explored its potential to edit adenosines within start codons (typically AUG), enabling translational interference. A challenge lies in the frequent presence of cytosine upstream of AUG^75^, which are suboptimal substrates for ADAR deaminases. To test this, we designed a reporter mRNA encoding SYFP2 with a start codon flanked by 5′ cytosine (Extended Data Fig. 11a). Remarkably, RECODEv2 induced a conversion rate of 53.3% (Extended Data Fig. 11b), demonstrating broad compatibility with previously unfavorable sites. To confirm the role of RNA editing on translational interference, we deleted Pepper from targeting gRNA to avoid ADAR1d recruitment. While minor editing (16.3%) occurred in the absence of Pepper, the stronger SYFP2 signal and similar RNA expression levels suggested that RNA editing played a substantial role in such process (Extended Data Fig. 11c, and 11d).

Finally, we applied RECODE to target the start codon of endogenous *SOD1* (Extended Data Fig. 11e). Silencing toxic *SOD1* mutants has been shown to mitigate ALS-related pathology^76^. As expected, RECODEv1 achieved efficient editing (48.3%) and translational interference (51.4% reduction in protein and insignificant change in mRNA levels compared to NT controls) (Extended Data Fig. 11f-11h), highlighting its therapeutic promise as a translational interference tool.

## Discussion

This study presents RECODE, a novel strategy for achieving precise and efficient RNA editing. Built upon the UDeg-Pepper system developed here, RECODE utilizes conditional ADAR1d variants with gRNA-regulated stability. This innovation enables degradation of off-target ADAR1d, while maintaining robust on-target activity. Notably, RECODE also displays reduced bystander editing, which can likely be attributed to the shortened guide-target hybrid length and the controlled abundance of ADAR1d. As a result, RECODE demonstrates superior specificity and efficiency compared to existing RNA editing approaches, representing a substantial advancement in the field. Moreover, the inherent modularity of RECODE facilitates integration with other naturally occurring or engineered deaminases^31, 38^, either to further enhance specificity or to expand the editing scope.

Despite these advancements, achieving the ideal scenario of complete target RNA editing with negligible off-target effects remains a challenging goal in the near future. A balance between activity and specificity is often necessary. Correspondingly, we developed two distinct versions of RECODE, each tailored for specific tasks. RECODEv1 gRNA recruits and stabilizes the deaminase with dual Peppers flanking the targeting sequence, significantly boosting editing yields, including bystander edits. RECODEv2 utilizes switchable gRNA with a locked Pepper design. This design, with carefully calibrated locking strength, allows switching of Pepper from “off” to “on” states upon guide-target binding, thereby minimizing the stabilization of off-target deaminases. The development of RECODEv2 introduces a novel mechanism for precise control of deaminase level through gRNA engineering. It enables the creation of RNA base editors with distinct specificity and activity by simply fine-tuning the locking stem of the gRNA. Identification of gRNA thermodynamic parameters that correlate with ADAR1d expression level and editing efficiency further streamlines this process.

Furthermore, by linking guide-target binding with protein stability, this mechanism holds potential for applications in programmable RNA sensing^61^. Considering the strong association between RNA signatures and cellular identities^77^, programmable RNA sensing presents a promising approach for targeting specific cell types and states. Our study contributes to this field by providing versatile UDegs, as well as guidelines for gRNA design.

Additionally, the development of UDegs carries significant implications for the rational design of potent and miniature protein destabilization domains. Such domains consist of two types of fundamental units: E3 ubiquitin ligase recognition motifs and preferred ubiquitination sites, with each unit comprising minimal amino acid residues. By identifying E3 ubiquitin ligase recognition motifs^45, 46^, it is anticipated that more efficient and custom protein degradation domains can be designed following the principles outlined here.

In parallel with efforts to minimize off-target effects, we identified hyperactive ADAR1d mutants using a structure-guided consensus approach^78^. This protein engineering strategy, inspired by natural evolution, achieved high success rates with minimal screening requirements. Moreover, we provide structural insights into how these modifications may enhance activity, using a model generated by Alphafold3^49^. These insights could further guide protein design. This accurate structural prediction using machine learning, combined with natural evolution-informed mutagenesis, highlights a powerful methodology for advancing protein engineering. Therefore, we envision that the principles underlying RECODE could be broadly applicable across diverse applications.

---

**Extended Data Fig. 1.**
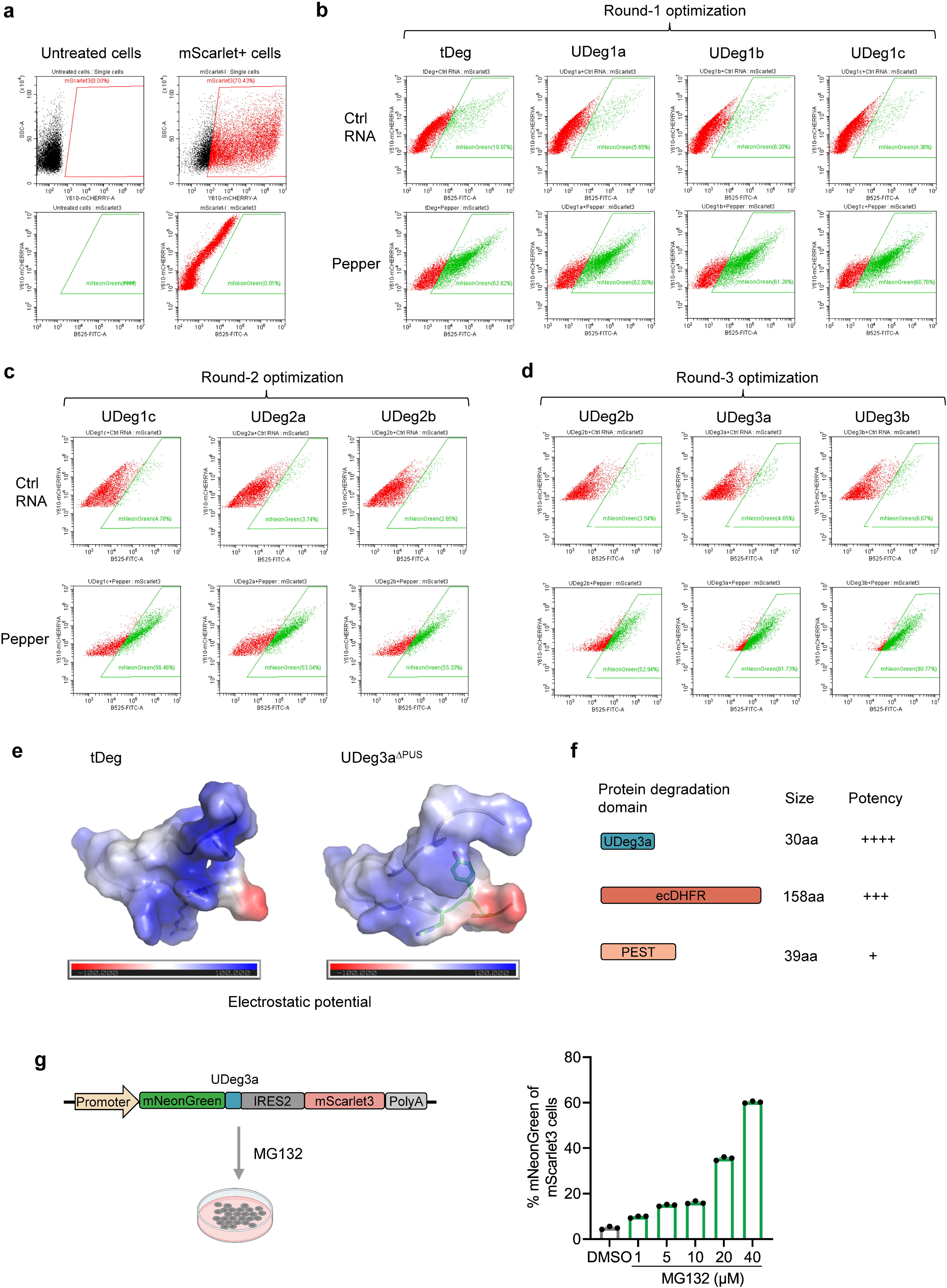
Development and characterization of UDeg3a. **a**, Representative plots for untreated and mScarlet3 single positive controls. **b**–**d**, Representative plots showing three rounds of optimization of tDeg, generating UDeg1 (**b**), UDeg2 (**c**), and UDeg3 (**d**). **e**, Electrostatic potential of tDeg and of UDeg3a with the PUS removed **f**, Size and strength comparison of the protein degradation domain: UDeg3a, ecDHFR, and PEST. **g** UPS-dependent degradation of UDeg3a. Left, schematic overview of the experimental design. Right, dose-dependent stabilization of UDeg3a by MG132, a proteosome inhibitor. Data are presented as mean ± SEM from 3 technical repeats.

**Extended Data Fig. 2.**
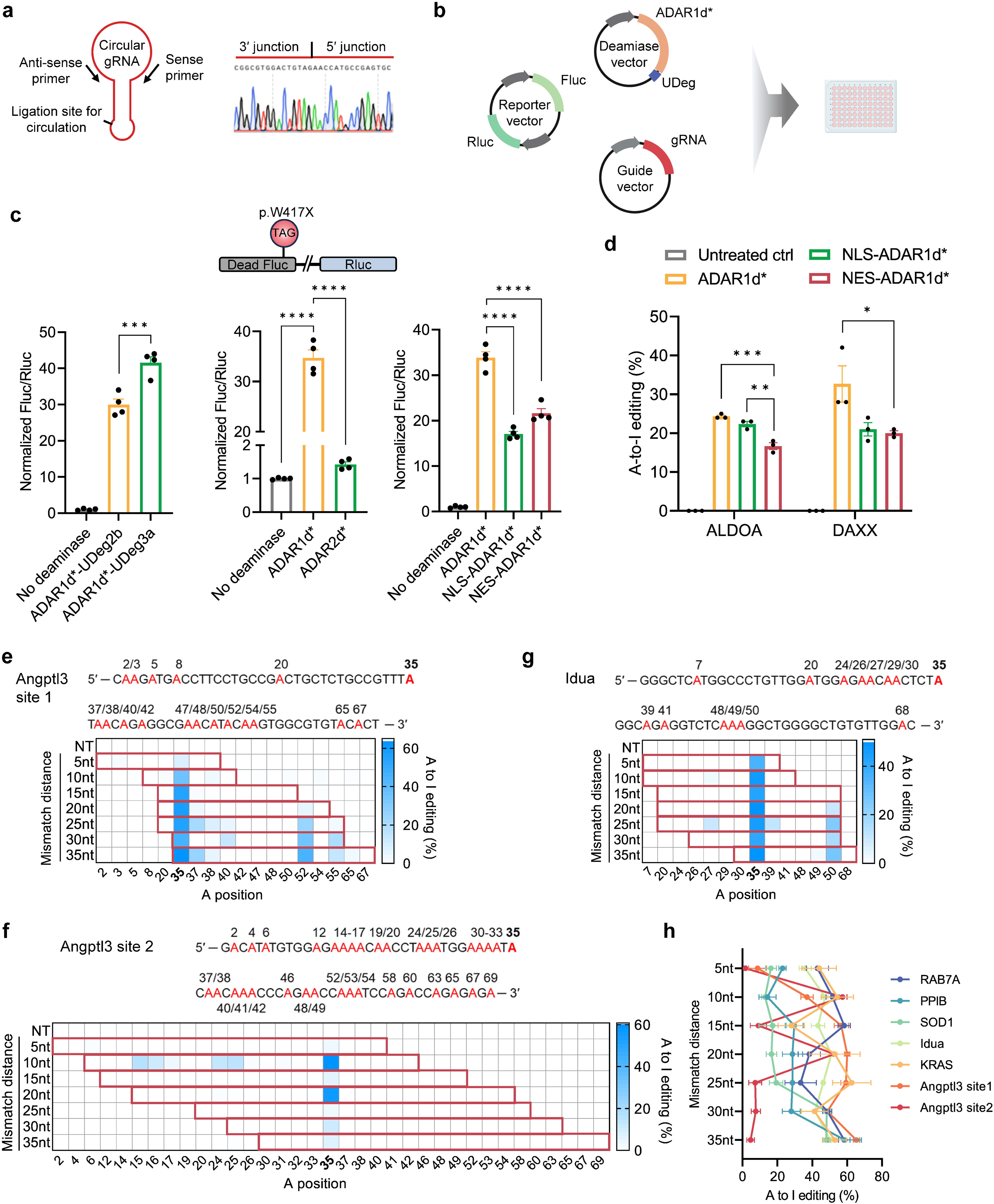
Development and characterization of RECODEv1. **a**, Circulation of gRNA validated by Sanger sequencing of PCR amplicons spanning the ligation site. **b**, Schematic overview of dual luciferase assays for assessing RNA editing efficiency. **c**, Deaminase-dependent RNA editing. (Left) increased binding between ADAR1d* and Pepper enhanced RNA editing activity. (Middle) When coupled with the UDeg3a-Pepper system, ADAR1d* is much more efficient than ADAR2d* in inducing RNA edits. (Right) ADAR1d*-UDeg3a is most efficient without specific localization signals. Data are presented as mean ± SEM from 4 biological repeats. One-way ANOVA: \*\*\**p*<0.001, **** *p*<0.0001. **d**, Editing of endogenous RNA confirms the optimal activity of ADAR1d*-UDeg3a without specific localization signals. Data are presented as mean ± SEM from 3 biological repeats. One-way ANOVA: * *p*<0.05, ** *p*<0.01\*\*\**p*<0.001. **e**–**g**, Heatmaps showing the editing rates of all adenosines, highlighted in red, covered by guides with variable mismatch distances targeting transfected Angptl3 (**e**,**f**) and Idua (**g**) transcripts. The target adenosines are emphasized in bold. For each guide, the covered adenosines are outlined in red in the heatmaps. Angled numbers below heatmaps indicate nucleotide positions from 1 to 69. **h**, The effect of mismatch positions on the editing efficiency of target adenosines. Data are presented as mean ± SEM from 3∼4 biological repeats.

**Extended Data Fig. 3.**
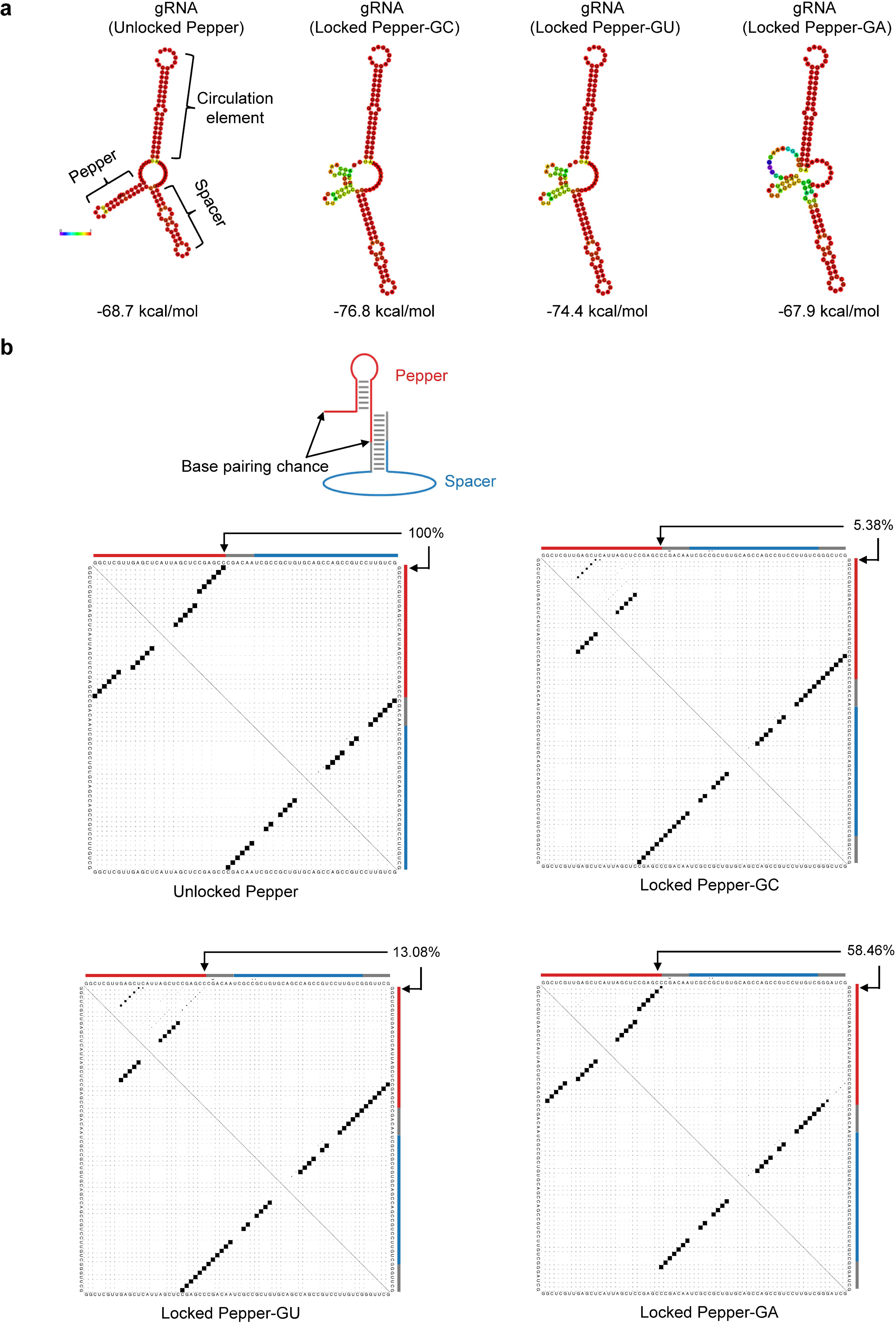
Thermodynamic analysis of gRNA folding. **a**, Predicted secondary structures of gRNAs with minimum free energy. The Pepper (recruitment domain) and spacer (targeting domain), and circulation domain are illustrated in gRNA of unlocked Pepper. Minimum and maximum base pairing probabilities are displayed by blue and red colors, respectively. The minimum free energies are labeled below the graphics. **b**, Partition function of gRNA folding. gRNA, excluding the universal circulation domain, was subject to folding prediction. Pepper folding probability is defined by the pairing likelihood of terminal base pair of Pepper. (Top) Graphic illustrating the locked Pepper and the disrupted terminal base pair of Pepper. (Bottom) Dot plots showing base pairing probabilities for each possible pair of bases. The terminal base pair of Pepper and its pairing likelihood are marked by arrows for each gRNA. Nucleotides constituting Pepper or spacer are distinguished by the colored lines.

**Extended Data Fig. 4.**
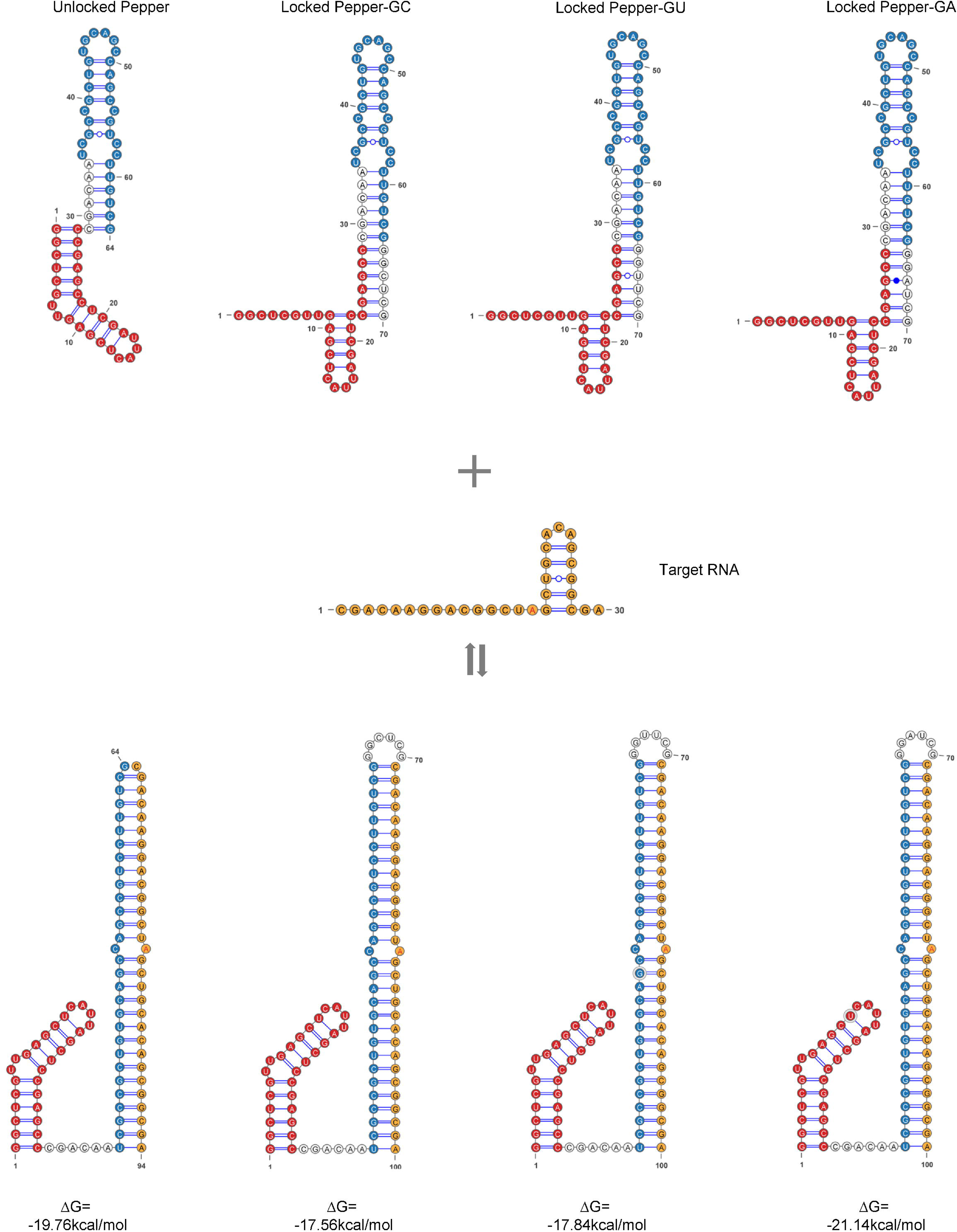
gRNA co-fold with target. Schematic showing the secondary structures of gRNA and target, and their dimerization. Delta G for heterodimerization is shown at the bottom. The circulation elements of gRNA are not shown here. Pepper, spacer, and target are marked by red, blue, and yellow filling colors, respectively. The target adenosine is highlighted in red.

**Extended Data Fig. 5.**
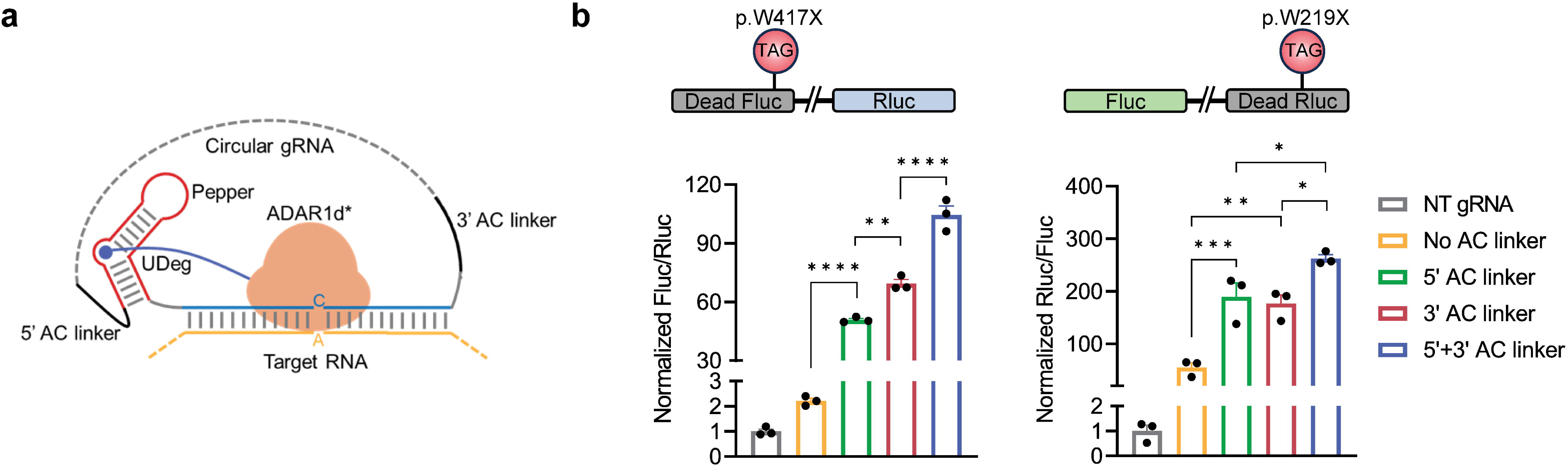
The effects of flanking AC linkers on gRNA performance. **a**, Schematic showing the positions of AC linkers in gRNA **b**, Luciferase assays revealed that appending AC linkers to the 5’ terminus of Pepper and the 3’ terminus of the spacer increased editing efficiency. Data are presented as mean ± SEM from 3 biological repeats. One-way ANOVA: \**p*<0.05, \*\**p*<0.01, \*\*\**p*<0.001, \*\*\*\**p*<0.0001.

**Extended Data Fig. 6.**
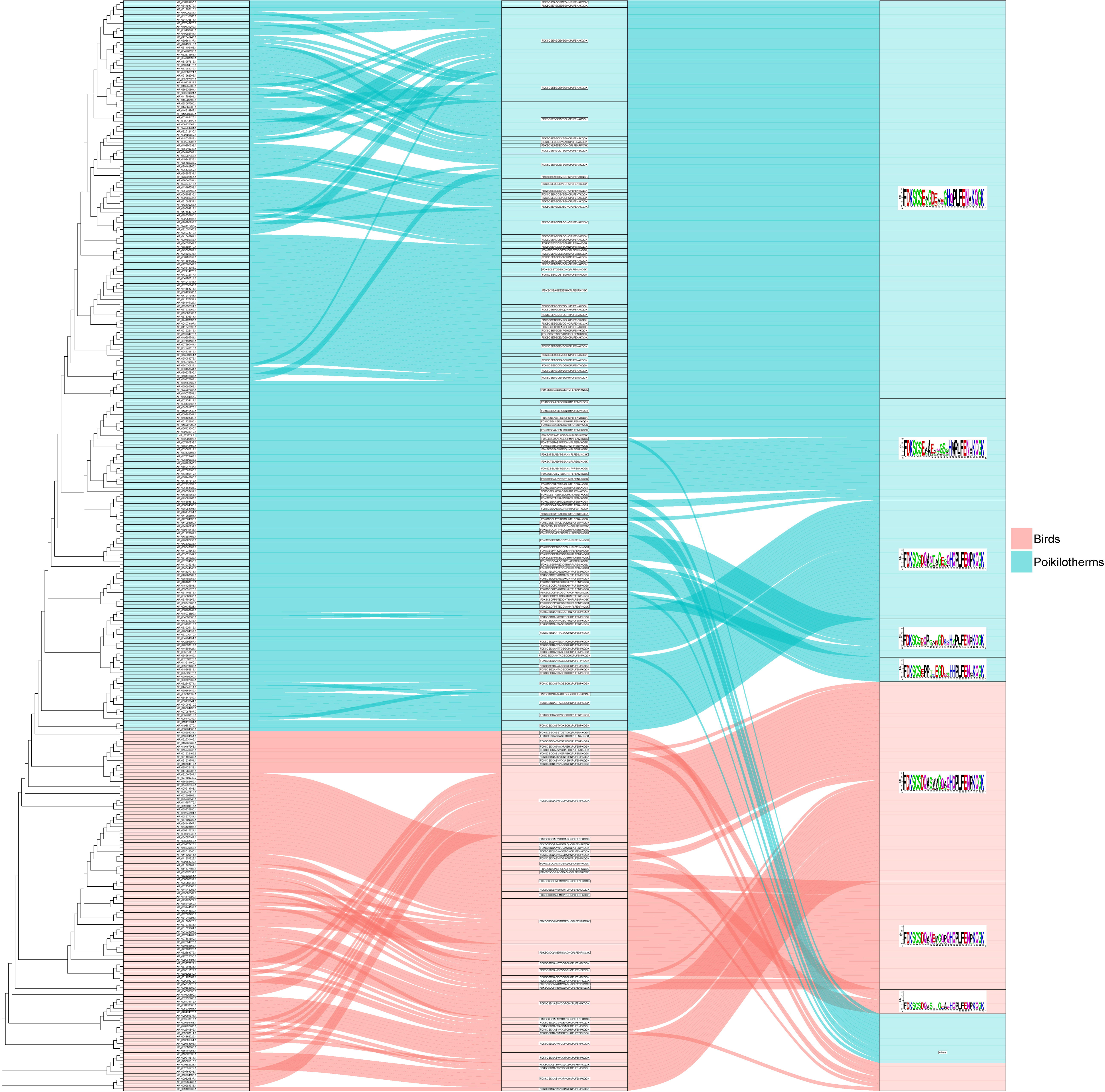
Sankey diagram illustrating the generation of consensus sequences of the ADAR1 5′ binding loop. Left node: phylogenetic tree of ADAR1 from poikilotherm vertebrates and Aves. Middle node: sequences of the 5′ binding loop. Right node: consensus of the 5′ binding loop.

**Extended Data Fig. 7.**
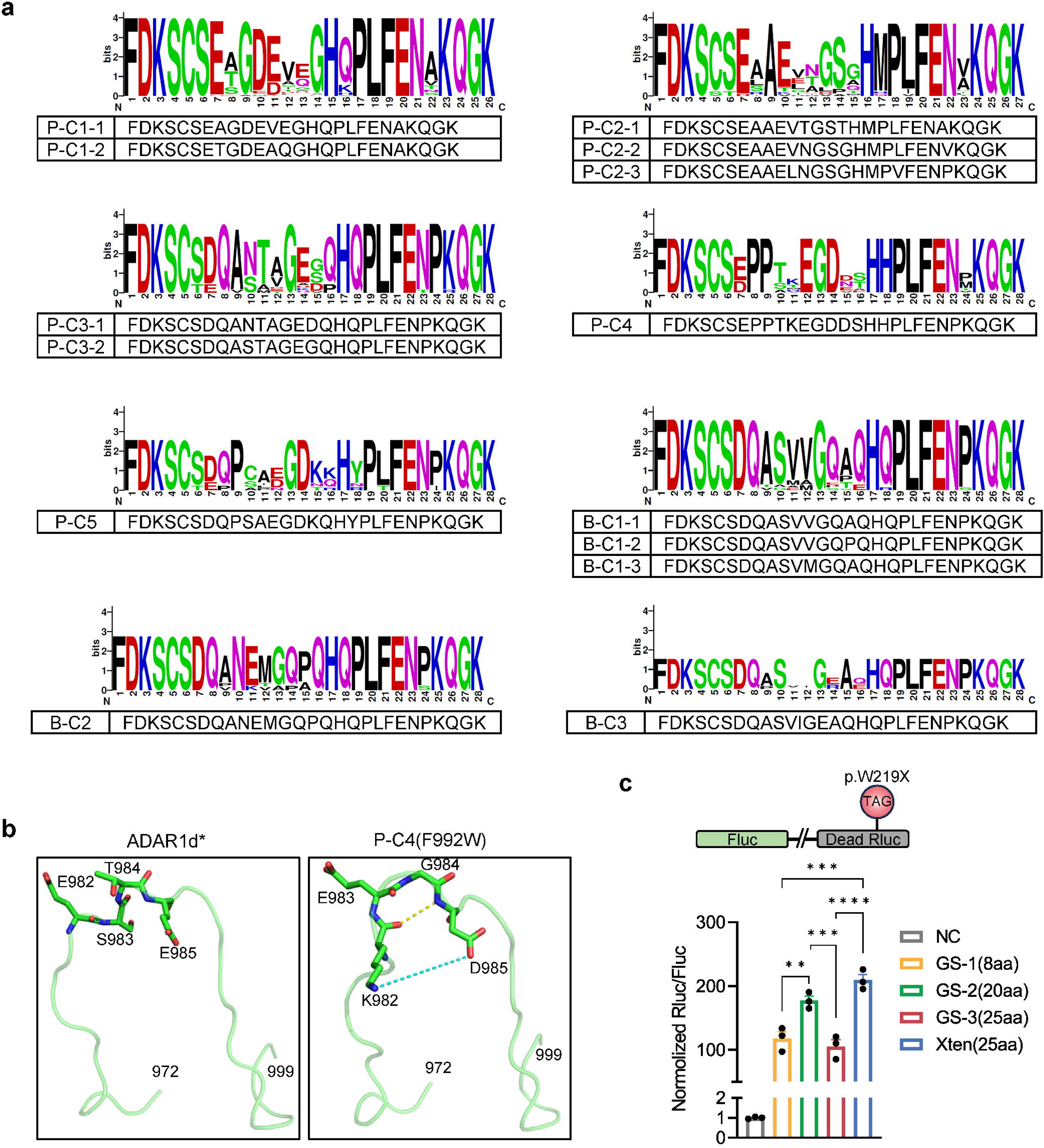
Engineering ADAR1d for improved activity. **a**, Consensus sequences of the ADAR1d 5′ binding loop from 8 clusters and the representative sequences of each motif. **b**, Structural analysis of the 5′ binding loop in ADAR1d and P-C4(F992W). In P-C4(F992W), the four amino acids (K982, E983, G984, D985) formed a classical β-turn at the loop apex, which is facilitated by electrostatic interactions (blue dashed lines) between K982 and D985. **c**, Linker optimization between P-C4(F992W) and UDeg3a. Data are presented as mean ± SEM from 3 biological repeats. One-way ANOVA: \*\**p*<0.01, \*\*\**p*<0.001, \*\*\*\**p*<0.0001.

**Extended Data Fig. 8.**
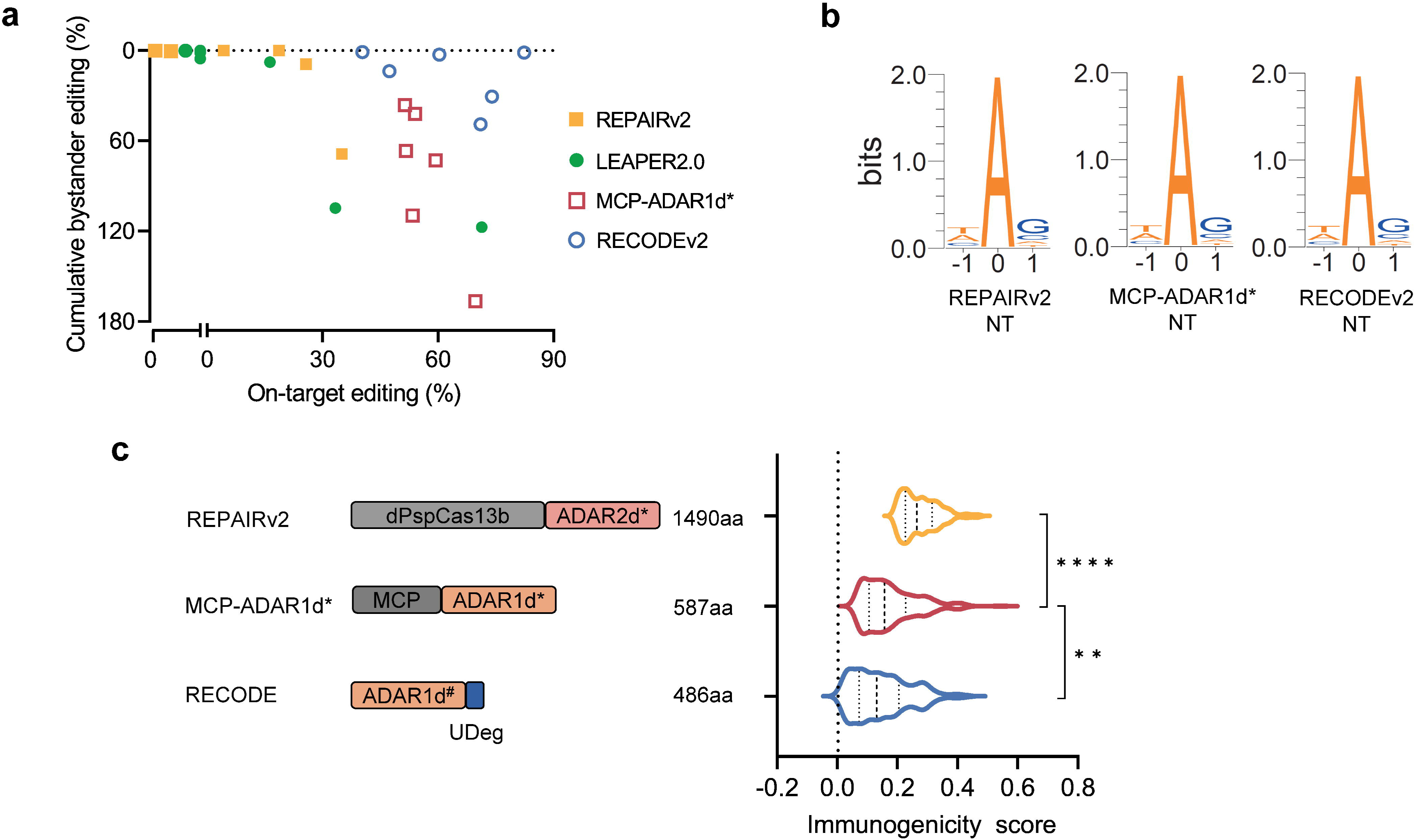
Comparing RECODE with established RNA editing methods. **a**, On-target versus cumulative bystander editing rates summarized from data in (Figure 5A-5F). **b**, Sequence logos generated from all A-to-I edits in the transcriptomes of HEK293T cells treated by non-targeting REPAIRv2, MCP-ADAR1d*, and RECODEv2. **c**, Schematic size comparison (left) of protein components from REPAIRv2, MCP-ADAR1d*, and RECODEv2. ADAR1d# denotes P-C4(F992W). Violin plots (right) of immunogenicity scores of peptides from REPAIRv2, MCP-ADAR1d*, and RECODEv2. Top 200 peptides ranked in predicted immunogenicity are subjected to analysis for each protein. Solid lines represent the median, and dashed lines denote the first and third quantiles within each violin. One-way ANOVA: ** *p*<0.01, \*\*\*\**p*<0.0001.

**Extended Data Fig. 9.**
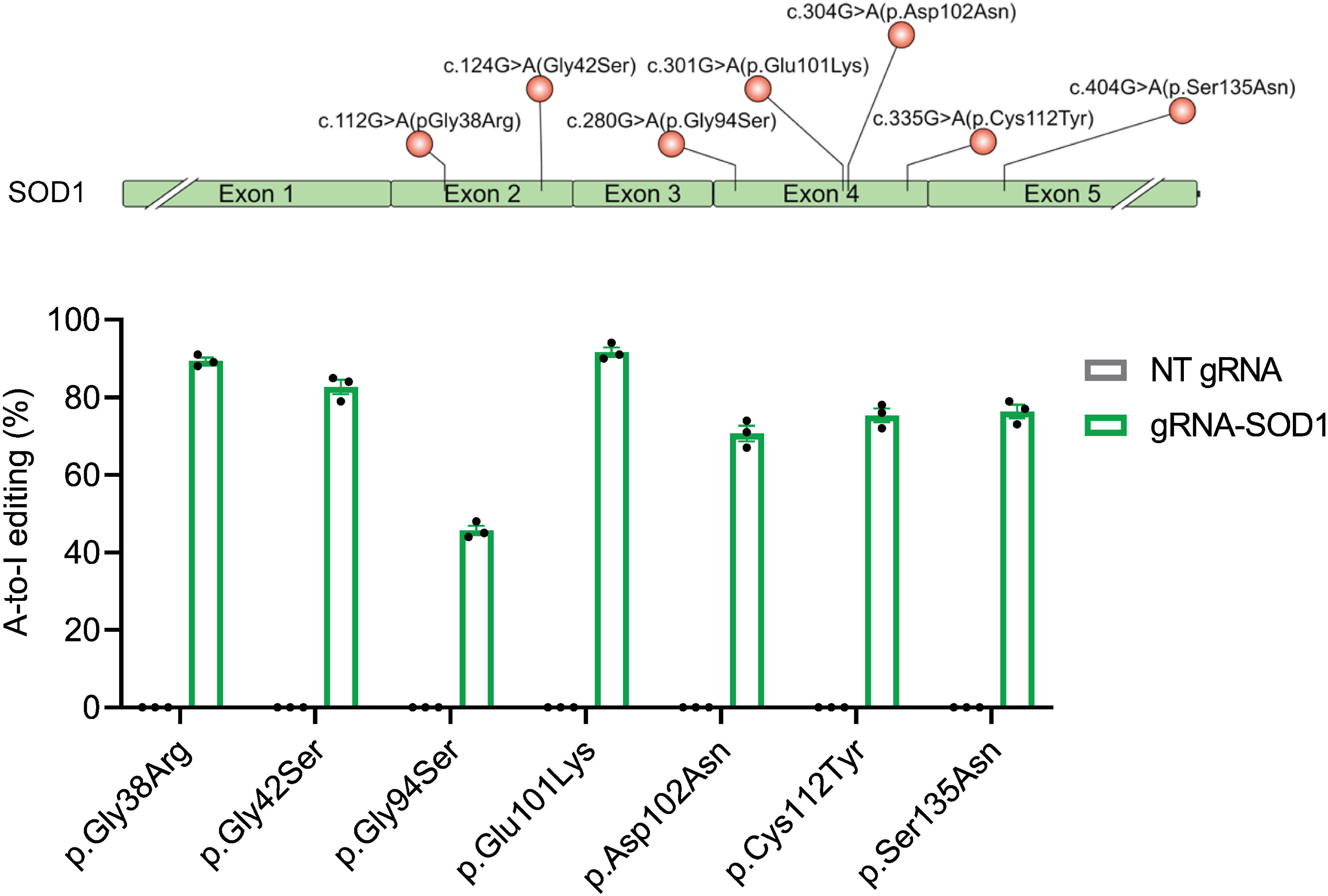
Robust editing of multiple ALS-relevant mutations in *SOD1* by RECODEv2. (Top) Schematic showing G-to-A mutations in the *SOD1* gene, targeted by RECODEv2. (Bottom) Quantification of the conversion rates. Data are presented as mean ± SEM from 3 biological repeats.

**Extended Data Fig. 10.**
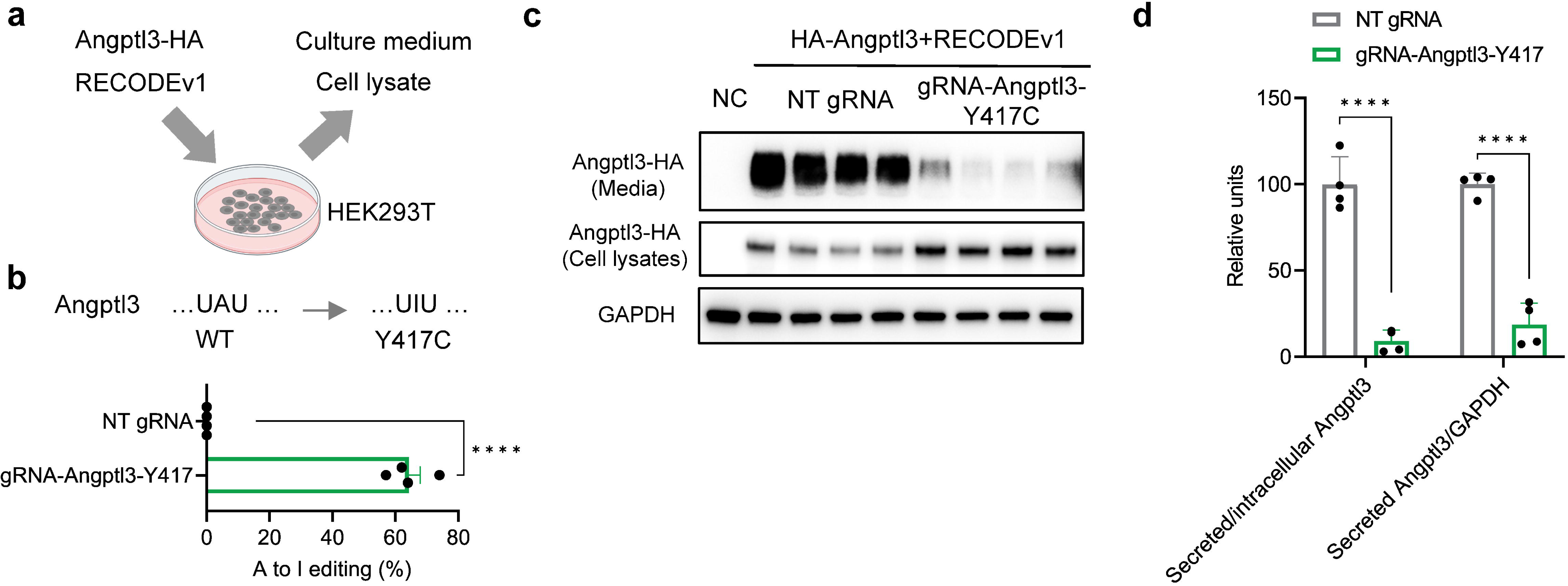
Installing a Y417C mutation into Angptl3 suppressed extracellular secretion by RECODEv1. **a**, Schematic showing the installation of a protective mutation into Angptl3 mRNA and assessment of protein secretion to culture medium. **b**, Histogram showing conversion rates of codon UAU to UIU. **c**, Western blot analysis of intracellular and secreted Angptl3-HA. **d**, Quantitative analysis of the ratios of secreted Angptl3-HA to either intracellular Angptl3-HA or GAPDH. For **b**,**d**, data are presented as mean ± SEM from 4 biological repeats. Two-tailed unpaired *t*-test: \*\*\*\**p*<0.0001.

**Extended Data Fig. 11.**
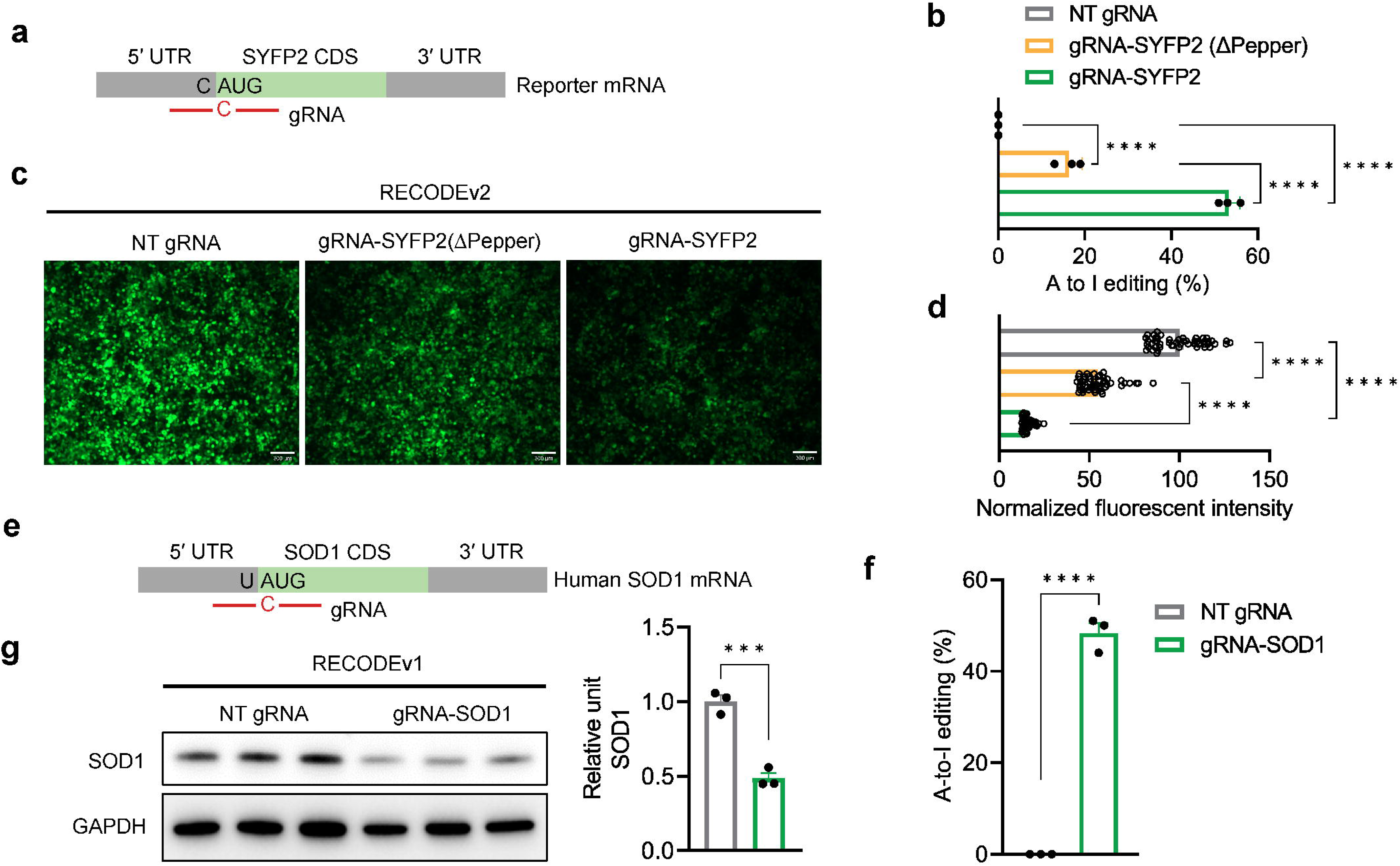
Harnessing RECODE for translational interference via start codon targeting. **a**, Schematic showing gRNA targeting start codon of reporter mRNA. **b**, Histograms showing the editing efficiency of adenosine within SYFP2 start codon by RECODEv2. Data are presented as mean ± SEM from 3 biological repeats. One-way ANOVA: **** *p*<0.0001. **c**, qPCR analysis of SYFP2 transcripts targeted by RECODEv2. Data are presented as mean ± SEM from 3 biological repeats. One-way ANOVA: \*\*\*\**p*<0.0001. **d**, Representative images and quantitative analysis of SYFP2 fluorescence in HEK293T cells following treatment by RECODEv2 NT gRNA, targeting gRNA lacking Pepper, and intact targeting gRNA. Scale bar, 300 μm. Data are presented as mean ± SEM from n=51 cells per group. One-way ANOVA: **** *p*<0.0001. **e**, Schematic of gRNA targeting the start codon of endogenous SOD1 mRNA. **f**, Histogram showing the editing efficiency of adenosine within SOD1 start codon. **g**, qPCR analysis of *SOD1* transcripts targeted by RECODEv1. Data are presented as mean ± SEM from 3 biological repeats. **h**, Western blot analysis (top) of SOD1 in HEK293T cells following treatment by RECODEv1. Quantitative analysis (bottom) of immunoblot bands reveals significant translational interference by RECODEv1. For **f**–**h**, data are presented as mean ± SEM from 3 biological repeats. Two-tailed unpaired *t*-test: ** *p*<0.01, **** *p*<0.0001.

## Methods

### Plasmid construction

Fluorescent protein reporter plasmids were generated by inserting mNeonGreen-IRES2-mScarlet3, synthesized by GENEWIZ, into pCDNA3.1(+) (Thermo). tDeg variants were then fused to the C terminus of mNeonGreen.

For the deaminase vector, the ADAR1 deaminase^E1008Q^ gene was cloned from MS2_adRNA-MCP_ADAR1 DD (E1008Q)_NES (Addgene) and inserted into pCI (Promega) for mammalian cell expression. Oligonucleotides encoding the consensus 5′ binding loop were synthesized by GENEWIZ and used to generate ADAR1 deaminase mutants via seamless cloning. For the guide vector, gRNA backbones carrying Pepper, preceded by U6 promoter were synthesized by GENEWIZ and cloned to pUC19 (Addgene) vector. Then the guide vector was modified accordingly to target distinct sites. The gRNA sequences are provided in Supplemental Table S1. The luciferase genes were cloned from pMirGlo (Promega) and subject to site-directed mutagenesis to induce non-sense mutations, followed by insertion into pCI (Promega).

Human FUS and SOD1 genes were amplified from cDNA of Hela cells and mouse Idua and Angptl3 genes were amplified from cDNA mouse livers, followed by the introduction of the mutations via site-directed mutagenesis. Then they were cloned into pCI for mammalian cell expression. FUS was further fused by SYFP2 or a HA tag at the N terminus.

The ADAR1 deaminase gene and U6-driven gRNA were assembled into pAAV-hSyn-EGFP (Addgene, #50465) to generate AAV vectors.

### AAV production and titration

For AAV production, HEK293T cells were co-transfected with a mixture of plasmids: an AAV vector plasmid, a packaging plasmid encoding the AAV2 Rep protein and capsid proteins for AAV9, and the pHGT1-Adeno1 helper plasmid carrying adenoviral genes. Transfections were conducted using calcium phosphate when HEK293T cells reached approximately 80% confluence in two 15-cm culture dishes. After 48–72 hours, the cells were harvested and resuspended in a buffer containing 150 mM NaCl and 100 mM Tris-HCl (pH 8.0). The cells were lysed through repeated freeze-thaw cycles, followed by the addition of MgCl2 to achieve a final concentration of 1 mM. AAV particles were then purified and concentrated using Millipore Amicon 100K columns (catalog no. UFC910008; Merck Millipore). Encapsidated viral DNA was quantified using qPCR (Thermo Fisher, Waltham, USA) with primers targeting the ITR sequences, following denaturation of the AAV particles with Proteinase K. Titers are expressed as viral genomes per milliliter (VG/mL).

### Cell culture and transfection

HEK293T, Huh7, and HepaG2 cells were cultured in Dulbecco’s Modified Eagle’s Medium (Thermo Fisher Scientific) supplemented with 10% fetal bovine serum (10091148, Gibco) and 1% penicillin-streptomycin (15140122, Gibco). The cells were maintained at 37 °C in a 5% CO2 atmosphere and regularly passaged using 0.05% Trypsin-EDTA (40127ES60, Yeasen).

Transfections were performed when the cells reached 60–80% confluence in 24-well plate or 96-well plate (Corning). All plasmids were delivered by jetOPTIMUS transfection reagent (101000006, Polyplus), according to the manufacturer’s protocol. The mass of plasmids used is specified for each experiment.

### Assessment of protein degradation domains

For UDeg engineering, HEK293T cells were seeded in 24-well plates, 1 day prior to transfection. of Reporter plasmids (200 ng) and Pepper-encoded plasmids (300 ng) were co-delivered into each well. For comparison between degrons, 500 ng, 1 μg, and 1.5 μg of reporter plasmids were used for HEK293T, Huh7, and HepaG2 cells, respectively. At 48 hours post transfection, cells were rinsed once with 1X DPBS (Servicebio) and dissociated with Trypsin (Yeasen). Cells were then collected and resuspended in 1X PBS, before flow cytometry analysis was performed.

### Flow cytometry analysis

Cells were resuspended in 1X PBS on ice. Live cells were gated by side scatter area versus forward scatter area (SSC-A vs. FSCA), singlets were selected by forward scatter height versus forward scatter area (FSC-H vs. FSC-A), and the fluorescence-positive population was gated against the mock-transfected control.

### Luciferase assays

HEK293T cells were seeded in 96-well polystyrene cell culture microplates (No. 655098, Greiner). Cells were transfected by a mixture of four plasmids (100 ng of deaminase vector, 25 ng of guide vector, 25 ng of dead luciferase vector, and 10 ng of another intact luciferase vector as an internal control). Each experimental group was set up with four replicate wells. After 48 hours, Firefly and Renilla luciferase fluorescence values were measured separately using Dual Glo Luciferase Reporter Gene Assay Kit (11405ES80, Yeasen), according to the manufacturer’s protocol. Bioluminescence was calculated as the ratio of Firefly to Renilla luciferase activity, or vice versa.

### RNA editing rate assessment and qPCR

For endogenous transcript editing, a total of 1.2 μg plasmids were used for each method, with a ratio of 3:1 for deaminase vectors to guide vectors. The ratio was adjusted to 6:2:1 (Deaminase:Guide:Target) in the cases of editing transfected transcripts. At 48 hours post-transfection, cells were collected. Total RNA was extracted using RNAiso Plus (9109, Takara) according to the manufacturer’s protocol. RNA was quantified using a NanoDrop spectrophotometer, and reverse transcription was performed using the RevertAid First Strand cDNA Synthesis Kit (Thermo Scientific) according to the kit instructions. qPCR was carried out with Hieff UNICON® Advanced qPCR SYBR Master Mix (Yeasen). The primer sequences were shown in Supplemental Table S2.

To evaluate the editing efficiency, PCR amplicons spanning editing sites were sent to Tsingke Biotechnology for Sanger sequencing. A-to-I conversion rates were quantified using the online tool editR (https://moriaritylab.shinyapps.io/editr_v10/).

### Western blot

Cells were harvested 72 hours after transfection and lysed in RIPA Lysis Buffer (P0013C, Beyotime) supplemented with Protease Inhibitor Cocktail (11836153001, Roche). Following centrifugation, the supernatant was collected as the protein sample. Protein concentration was measured using the Pierce BCA Protein Assay Kit (23225, Thermo Scientific), after which the samples were mixed with 5×SDS-PAGE Protein Loading Buffer (G2075, Servicebio) and denatured by heating.

Protein separation was performed via SDS-PAGE electrophoresis according to standard procedures, followed by transfer to 0.45-µm polyvinylidene fluoride (PVDF) membranes (Merck Millipore).

Membranes were incubated with the appropriate primary antibody, followed by HRP-conjugated secondary antibody (APExBIO). Protein signals were visualized using the ECL Chemiluminescent Substrate Detection Kit (K1231, APExBIO) on a Tanon 5200 Chemiluminescent Imaging System.

The HA antibody was used for detection of ADAR1 deaminase. For immunoblotting of SOD1, nuclear and cytosolic fractions of cell lysate were prepared separately using a Nuclear and Cytoplasmic Protein Extraction Kit (P0027, Beyotime). Then, the proteins of interest were detected as described above.

Protein bands were quantified using ImageJ software to assess protein expression levels. The intensity of the band of interest was normalized by dividing it by the corresponding α-tubulin or GAPDH bands on the same membrane after stripping.

### AAV delivery in adult mice

All animal-related procedures were performed according to protocols approved by the Institutional Animal Care and Use Committee (IACUC) at the Shenzhen Institutes of Advanced Technology (SIAT), Chinese Academy of Sciences. All mice were maintained following standard housing conditions.

For unilateral AAV delivery to the M1 brain area of adult mice, stereotaxic injections were performed. The coordinates used for the M1 were: X (ML) +2.0mm, Y (AP) +2.1mm, and Z (DV) –1.7mm. Mice were anesthetized with isoflurane and placed in a stereotaxic frame (Reward Life Science). The surgical area was cleaned using 2% iodine and 70% alcohol. The scalp overlying the dorsal skull was incised and a small hole was drilled over the injection site. AAV vectors were injected into the target brain regions using a 10 µl Hamilton syringe, connected to a pump. After injection, the needle was left in place for 5 minutes before being slowly withdrawn. Once the needle was removed and the scalp sutured, the mouse was removed from the stereotaxic apparatus and kept on a heating pad until after movement recovery, at which point it was brought back to its home cage.

### Immunostaining

Brain tissues were collected and fixed in 4% paraformaldehyde at 4 °C overnight, followed by cryoprotection in 30% sucrose until the tissues sank. The tissues were then embedded in OCT and cryosectioned at a thickness of 20 µm. Sections were permeabilized with 0.3% Triton X-100 in PBS for 30 minutes, followed by blocking with 5% goat serum in PBS for 1 hour at room temperature. Primary antibodies were applied and incubated overnight at 4 °C. After washing with PBS, sections were incubated with secondary antibodies conjugated to fluorescent dyes for 1 hour at room temperature. Nuclei were stained with DAPI, and sections were mounted with antifade mounting medium. Images were captured using a fluorescence or confocal microscope.

For the imaging of SYFP2-FUS, cells were grown on glass slides. At 48 hours post-transfection, cells were fixed in 4% paraformaldehyde for 10 minutes at room temperature. Then, the cells were mounted with mounting medium and DAPI for confocal imaging.

### Multiple sequence alignment

Orthologous ADAR1 sequences of fish, reptiles, and birds were retrieved from NCBI and were aligned using the Clustal Omega. Poorly aligned regions were manually adjusted if required. Phylogenetic trees were generated with FastTree (Arkin Laboratory). Sequences of ADAR1 5’ binding loop were grouped by the Cluster tool in the Immune Epitope Database, followed by a sequence consensus for each group by WebLogo.

### Structure prediction by Alphafold3

Sequences of tDeg and UDeg3a excluding PUS, as well as that of Pepper, were input into Alphafold3 to analyze their interactions, with default parameters set. Similar procedures were also applied to ADAR1 deaminase. Then, the structures were visualized and annotated using PyMOL (Schrödinger).

### RNA folding analysis

The secondary structures of gRNA were predicted using ViennaRNA Web Services (http://rna.tbi.univie.ac.at/). The minimum free energy and partition function of gRNAs were analyzed using RNAfold. The free energy of gRNA binding to target was calculated using RNAcofold. The predicted secondary structures were visualized using VARNA software.

### Immunogenicity prediction

The protein sequences of dPspCas13b-ADAR2d, MCP-ADAR1d*, and P-C4(F992W)-UDeg3a were uploaded to Immune Epitope Database (IDEB) for prediction of immunogenicity using T cell class I tool. NetMHCpan EL 4.1 (IEDB recommended epitope predictor 2023.09) was used to predict the strength of the peptide:MHC interaction. Top 200 predicted MHC class I restricted 9-mer candidates were subjected to analysis for each protein.

### Transcriptome sequencing

HEK293T cells were transfected with 200 ng of plasmid DNA (Deaminase:Guide = 3:1) for each well in 24-well plates. Forty-eight hours post-transfection, total RNA was extracted using RNAiso Plus (Takara). The poly-A RNAs were enriched from total RNA using Dynabeads mRNA Purification Kit (Cat#61006, Invitrogen) and fragmented into small pieces using fragmentation reagent in MGIEasy RNA Library Prep Kit V3.1 (Cat# 1000005276, MGI). The first strand of cDNA was synthesized using random primes and reverse transcriptase, followed by second strand cDNA synthesis. The synthesized cDNA was end-repaired, A-tailing was added and then ligated to the sequencing adapters according to library construction protocol. The cDNA fragments were amplified by PCR and purified with MGIEasy DNA Clean beads (CAT#1000005279, MGI). The library was analyzed using an Agilent Technologies 2100 bioanalyzer. The double stranded PCR products were heat denatured and circularized by the splint oligo sequence in MGIEasy Circularization Module (CAT#1000005260, MGI). The single strand circle DNA (ssCir DNA) was formatted as the final library. The qualified libraries were sequenced using the DNBSEQ-T7RS platform.

### RNA-seq data analysis

To obtain clean data, the raw sequencing data were filtered using fastq (v0.23.4) with the following parameters: -q 15 -u 40 --detect_adapter_for_pe. The filtered clean data were then aligned to the GRCh38 reference genome using STAR (v2.7.11b). The alignment results were sorted with samtools. Next, PCR duplicates were removed using the gatk (v4.5.0.0) MarkDuplicates module (parameters: --CREATE_INDEX true --VALIDATION_STRINGENCY SILENT). The reads were split into exon segments, removing Ns while retaining grouping information, and sequences overhanging into intronic regions were removed using the gatk SplitNCigarReads module with default parameters.

Variant calling was performed with the gatk HaplotypeCaller module (parameters: -dont-use-soft-clipped-bases --standard-min-confidence-threshold-for-calling 20). The generated GVCF files were merged using the gatk CombineGVCFs module with default settings, followed by variant genotyping with the gatk GenotypeGVCFs module. SNP variants were selected using the gatk SelectVariant module (parameters: -select-type SNP). Filtering was applied using the gatk VariantFiltration module with the following criteria: --filter-expression “QD < 2.0 || MQ < 40.0 || FS > 60.0 || SOR > 3.0 || MQRankSum < -12.5 || ReadPosRankSum < -8.0”. For analyzing A-to-G variants, the proportion of G depth relative to the total depth in edited samples was calculated when control sample was AA.

### Statistical analysis

All statistics were performed using GraphPad Prism 9.0. The following tests were used as appropriate: paired *t*-test, unpaired *t*-test, one-way ANOVA, Tukey’s test, and Dunnett’s test. All *t*-tests were performed as two-tailed. In all statistical tests, a *p*-value <0.05 was considered statistically significant. Sample sizes were chosen based on previous publications or previous experience to yield sufficient power to detect specific effects. All statistical tests used are indicated in the figure legends.

## Data and code availability

All data reported in this paper will be shared by the lead contact upon reasonable request. No original code was generated in this study.

## Acknowledgements

We thank Dr. Chunshan Deng, Dr. Yinqing Li, and members of the Lu laboratory for helpful discussions and comments. This research was supported by Strategic Priority Research Program of the Chinese Academy of Sciences grant XDB0930000 (Z.L.), Shenzhen Medical Research Fund B2402029 (Z.L.) and B2302053 (Z.L.), National Natural Science Foundation of China grants 32400927 (Z.H.) and 82327805 (Z.L.), Shenzhen Science and Technology Innovation Commission grant KQTD20210811090117032 (Z.L.), China Postdoctoral Science Foundation grant 2023M743681 (Z.H.), and Guangdong Basic and Applied Research Foundation grant 2023A1515110864 (Z.H.).

## Author contributions

T.L. and Z.L. conceived the project. T.L., Yunping Lin, Q.L., W.L., Y.Z., W.C. and Yunyi Lin developed methods and performed experiments. T.L., Yunping Lin, Q.L., L.Y., and Z.H. analyzed data. Z.L. supervised the project. T.L., Yunping Lin, Z.H., and Z.L. wrote the manuscript. All authors read and approved the manuscript.

## References

1. Crick, F. Central Dogma of Molecular Biology. Nature 227, 561-& (1970).

2. Manning, K.S. & Cooper, T.A. The roles of RNA processing in translating genotype to phenotype. Nat Rev Mol Cell Bio 18, 102–114 (2017).

3. Moore, M.J. & Proudfoot, N.J. Pre-mRNA Processing Reaches Back to Transcription and Ahead to Translation. Cell 136, 688–700 (2009).

4. Wright, C.J., Smith, C.W.J. & Jiggins, C.D. Alternative splicing as a source of phenotypic diversity. Nat Rev Genet 23, 697–710 (2022).

5. Tian, B. & Manley, J.L. Alternative polyadenylation of mRNA precursors. Nat Rev Mol Cell Bio 18, 18–30 (2017).

6. Nishikura, K. Functions and Regulation of RNA Editing by ADAR Deaminases. Annu Rev Biochem 79, 321–349 (2010).

7. Eisenberg, E. & Levanon, E.Y. A-to-I RNA editing - immune protector and transcriptome diversifier. Nat Rev Genet 19, 473–490 (2018).

8. Liddicoat, B.J. et al. RNA editing by ADAR1 prevents MDA5 sensing of endogenous dsRNA as nonself. Science 349, 1115–1120 (2015).

9. Mannion, N.M. et al. The RNA-Editing Enzyme ADAR1 Controls Innate Immune Responses to RNA. Cell Rep 9, 1482–1494 (2014).

10. Pestal, K. et al. Isoforms of RNA-Editing Enzyme ADAR1 Independently Control Nucleic Acid Sensor MDA5-Driven Autoimmunity and Multi-organ Development. Immunity 43, 933–944 (2015).

11. Peng, X.X. et al. A-to-I RNA Editing Contributes to Proteomic Diversity in Cancer. Cancer Cell 33, 817-+ (2018).

12. Garrett, S. & Rosenthal, J.J.C. RNA Editing Underlies Temperature Adaptation in K^+^ Channels from Polar Octopuses. Science 335, 848–851 (2012).

13. Yablonovitch, A.L. et al. Regulation of gene expression and RNA editing in adapting to divergent microclimates. Nat Commun 8 (2017).

14. Pullirsch, D. & Jantsch, M.F. Proteome diversification by adenosine to inosine RNA-editing. Rna Biol 7, 205–212 (2010).

15. Dadush, A., et al. DNA and RNA base editors can correct the majority of pathogenic single nucleotide variants. Npj Genom Med 9 (2024).

16. Booth, B.J. et al. RNA editing: Expanding the potential of RNA therapeutics. Mol Ther 31, 1533–1549 (2023).

17. Qu, L. et al. Programmable RNA editing by recruiting endogenous ADAR using engineered RNAs. Nat Biotechnol 37, 1059–1069 (2019).

18. Katrekar, D. et al. Efficient in vitro and in vivo RNA editing via recruitment of endogenous ADARs using circular guide RNAs. Nat Biotechnol 40, 938–945 (2022).

19. Yi, Z.Y. et al. Engineered circular ADAR-recruiting RNAs increase the efficiency and fidelity of RNA editing in vitro and in vivo. Nat Biotechnol 40, 946-+ (2022).

20. Wettengel, J., Reautschnig, P., Geisler, S., Kahle, P.J. & Stafforst, T. Harnessing human ADAR2 for RNA repair - Recoding a PINK1 mutation rescues mitophagy. Nucleic Acids Res 45, 2797–2808 (2017).

21. Reautschnig, P. et al. CLUSTER guide RNAs enable precise and efficient RNA editing with endogenous ADAR enzymes in vivo. Nat Biotechnol 40, 759-+ (2022).

22. Merkle, T. et al. Precise RNA editing by recruiting endogenous ADARs with antisense oligonucleotides. Nat Biotechnol 37, 133-+ (2019).

23. Reautschnig, P. et al. Precise in vivo RNA base editing with a wobble-enhanced circular CLUSTER guide RNA. Nat Biotechnol (2024).

24. Monian, P. et al. Endogenous ADAR-mediated RNA editing in non-human primates using stereopure chemically modified oligonucleotides. Nat Biotechnol 40, 1093-+ (2022).

25. Yi, Z.Y. et al. Utilizing AAV-mediated LEAPER 2.0 for programmable RNA editing in non-human primates and nonsense mutation correction in humanized Hurler syndrome mice. Genome Biol 24 (2023).

26. Wang, Y.R., Park, S. & Beal, P.A. Selective Recognition of RNA Substrates by ADAR Deaminase Domains. Biochemistry-Us 57, 1640–1651 (2018).

27. Zambrano-Mila, M.S. et al. Dissecting the basis for differential substrate specificity of ADAR1 and ADAR2. Nat Commun 14 (2023).

28. Uhlen, M. et al. Tissue-based map of the human proteome. Science 347 (2015).

29. Tan, M.H. et al. Dynamic landscape and regulation of RNA editing in mammals. Nature 550, 249-+ (2017).

30. Abudayyeh, O.O. et al. A cytosine deaminase for programmable single-base RNA editing. Science 365, 382-+ (2019).

31. Huang, X.X. et al. Programmable C-to-U RNA editing using the human APOBEC3A deaminase. Embo J 39 (2020).

32. Buchumenski, I., et al. Global quantification exposes abundant low-level off-target activity by base editors. Genome Res 31, 2354–2361 (2021).

33. Katrekar, D. et al. In vivo RNA editing of point mutations via RNA-guided adenosine deaminases. Nat Methods 16, 239-+ (2019).

34. Cox, D.B.T. et al. RNA editing with CRISPR-Cas13. Science 358, 1019–1027 (2017).

35. Marina, R.J., Brannan, K.W., Dong, K.D., Yee, B.A. & Yeo, G.W. Evaluation of Engineered CRISPR-Cas-Mediated Systems for Site-Specific RNA Editing. Cell Rep 33 (2020).

36. Xu, C.L. et al. Programmable RNA editing with compact CRISPR-Cas13 systems from uncultivated microbes. Nat Methods 18, 499-+ (2021).

37. Liu, Y.J., et al. REPAIRx, a specific yet highly efficient programmable A &gt; I RNA base editor. Embo J 39 (2020).

38. Yan, H. & Tang, W.X. Programmed RNA editing with an evolved bacterial adenosine deaminase. Nat Chem Biol 20 (2024).

39. Wu, J.H. et al. Live imaging of mRNA using RNA-stabilized fluorogenic proteins. Nat Methods 16, 862-+ (2019).

40. Shaner, N.C. et al. A bright monomeric green fluorescent protein derived from. Nat Methods 10, 407-+ (2013).

41. Gadella, T.W.J. et al. mScarlet3: a brilliant and fast-maturing red fluorescent protein. Nat Methods 20, 541-+ (2023).

42. Guharoy, M., Bhowmick, P., Sallam, M. & Tompa, P. Tripartite degrons confer diversity and specificity on regulated protein degradation in the ubiquitin-proteasome system. Nat Commun 7 (2016).

43. Crowe, C. et al. Mechanism of degrader-targeted protein ubiquitinability. Science Advances 10, eado6492 (2024).

44. Fu, H.L., Yang, Y.X., Wang, X.B., Wang, H. & Xu, Y. DeepUbi: a deep learning framework for prediction of ubiquitination sites in proteins. Bmc Bioinformatics 20 (2019).

45. Yeh, C.W. et al. The C-degron pathway eliminates mislocalized proteins and products of deubiquitinating enzymes. Embo J 40 (2021).

46. Koren, I. et al. The Eukaryotic Proteome Is Shaped by E3 Ubiquitin Ligases Targeting C-Terminal Degrons. Cell 173, 1622-+ (2018).

47. Smith, C.A., Calabro, V. & Frankel, A.D. An RNA-Binding chameleon. Mol Cell 6, 1067–1076 (2000).

48. Smith, C.A., Crotty, S., Harada, Y. & Frankel, A.D. Altering the context of an RNA bulge switches the binding specificities of two viral Tat proteins. Biochemistry-Us 37, 10808–10814 (1998).

49. Abramson, J. et al. Accurate structure prediction of biomolecular interactions with AlphaFold 3. Nature 630 (2024).

50. Calabro, V., Daugherty, M.D. & Frankel, A.D. A single intermolecular contact mediates intramolecular stabilization of both RNA and protein. P Natl Acad Sci USA 102, 6849–6854 (2005).

51. Rogers, S., Wells, R. & Rechsteiner, M. Amino-Acid-Sequences Common to Rapidly Degraded Proteins - the Pest Hypothesis. Science 234, 364–368 (1986).

52. Iwamoto, M., Björklund, T., Lundberg, C., Kirik, D. & Wandless, T.J. A General Chemical Method to Regulate Protein Stability in the Mammalian Central Nervous System. Chem Biol 17, 981–988 (2010).

53. Lee, D.H. & Goldberg, A.L. Proteasome inhibitors: valuable new tools for cell biologists. Trends Cell Biol 8, 397–403 (1998).

54. Wang, Y.R., Havel, J. & Beal, P.A. A Phenotypic Screen for Functional Mutants of Human Adenosine Deaminase Acting on RNA 1. Acs Chem Biol 10, 2512–2519 (2015).

55. Litke, J.L. & Jaffrey, S.R. Highly efficient expression of circular RNA aptamers in cells using autocatalytic transcripts. Nat Biotechnol 37, 667-+ (2019).

56. Matthews, M.M. et al. Structures of human ADAR2 bound to dsRNA reveal base-flipping mechanism and basis for site selectivity. Nat Struct Mol Biol 23, 426–433 (2016).

57. Montiel-González, M.F., Vallecillo-Viejo, I.C. & Rosenthal, J.J.C. An efficient system for selectively altering genetic information within mRNAs. Nucleic Acids Res 44 (2016).

58. Wang, X., et al. Develop a Compact RNA Base Editor by Fusing ADAR with Engineered EcCas6e. Adv Sci 10 (2023).

59. Lehmann, K.A. & Bass, B.L. Double-stranded RNA adenosine deaminases ADAR1 and ADAR2 have overlapping specificities. Biochemistry-Us 39, 12875–12884 (2000).

60. Tyagi, S. & Kramer, F.R. Molecular beacons: Probes that fluoresce upon hybridization. Nat Biotechnol 14, 303–308 (1996).

61. Zhou, W.J. et al. Genetically Encoded Sensor Enables Endogenous RNA Imaging with Conformation-Switching Induced Fluorogenic Proteins. J Am Chem Soc 143, 14394–14401 (2021).

62. Hofacker, I.L. Vienna RNA secondary structure server. Nucleic Acids Res 31, 3429–3431 (2003).

63. Thomas, J.M. & Beal, P.A. How do ADARs bind RNA? New protein-RNA structures illuminate substrate recognition by the RNA editing ADARs. Bioessays 39 (2017).

64. Park, S. et al. High-throughput mutagenesis reveals unique structural features of human ADAR1. Nat Commun 11 (2020).

65. Yan, Z. et al. Next-generation IEDB tools: a platform for epitope prediction and analysis. Nucleic Acids Res 52, W526–W532 (2024).

66. Vance, C. et al. ALS mutant FUS disrupts nuclear localization and sequesters wild-type FUS within cytoplasmic stress granules. Hum Mol Genet 22, 2676–2688 (2013).

67. Shang, Y.L. & Huang, E.J. Mechanisms of FUS mutations in familial amyotrophic lateral sclerosis. Brain Res 1647, 65–78 (2016).

68. López-Erauskin, J. et al. ALS/FTD-Linked Mutation in FUS Suppresses Infra-axonal Protein Synthesis and Drives Disease Without Nuclear Loss-of-Function of FUS. Neuron 100, 1–15 (2018).

69. Wu, Z.J., Yang, H.Y. & Colosi, P. Effect of Genome Size on AAV Vector Packaging. Mol Ther 18, 80–86 (2010).

70. Braak, H. et al. Amyotrophic lateral sclerosis-a model of corticofugal axonal spread. Nat Rev Neurol 9, 708–714 (2013).

71. Wroe, R., Butler, A.W.L., Andersen, P.M., Powell, J.F. & Al-Chalabi, A. ALSOD: The Amyotrophic Lateral Sclerosis Online Database. Amyotroph Lateral Sc 9, 249–250 (2008).

72. Landrum, M.J. et al. ClinVar: public archive of interpretations of clinically relevant variants. Nucleic Acids Res 44, D862–D868 (2016).

73. Ono, M. et al. Protein region important for regulation of lipid metabolism in angiopoietin-like 3 (ANGPTL3) -: ANGPTL3 is cleaved and activated. J Biol Chem 278, 41804–41809 (2003).

74. Romeo, S. et al. Rare loss-of-function mutations in family members contribute to plasma triglyceride levels in humans. J Clin Invest 119, 70–79 (2009).

75. Kozak, M. An Analysis of 5’-Noncoding Sequences from 699 Vertebrate Messenger-Rnas. Nucleic Acids Res 15, 8125–8148 (1987).

76. Miller, T.M. et al. Trial of Antisense Oligonucleotide Tofersen for ALS. New Engl J Med 387, 1099–1110 (2022).

77. Snyder, M.P. et al. The human body at cellular resolution: the NIH Human Biomolecular Atlas Program. Nature 574, 187–192 (2019).

78. Porebski, B.T. & Buckle, A.M. Consensus protein design. Protein Eng Des Sel 29, 245–251 (2016).

